# Detection of diverse coronaviruses, paramyxoviruses, and rhabdoviruses from cave-dwelling bats in Eastern Uganda

**DOI:** 10.64898/2026.08.06.743307

**Authors:** Kayiwa John, Nassuna Charity, Nabatanzi Leonara, Yiga Fahim, Emma K Harris, Wickenkamp Natalie, Kalani M Williams, Matovu Benard, Mutebi Jack Micheal, Nalukenge Lillian, Nalikka Betty, Siya Aggrey, Nakayiki Teddy, Fagre Anna, Hartwick Anna, Cordova Emerald, Azerigyik Faustus, Castle Kevin, Dewey Tanya, Kityo N Robert, Lutwama J Julius, Rebekah C Kading

## Abstract

Bats harbor a diversity of viruses, some of which have the potential to impact human and livestock health. Caves in Eastern Uganda are commonly inhabited by bats in the genera *Rhinolophus, Hipposideros, Myonycteris,* and others. Human encroachment into these caves for shelter, hunting, mineral harvesting, and tourism poses a risk of exposure to infectious agents these bats may carry, yet little is known about the viruses present in these bats. From 2021 – 2023, 635 unique bats were captured in caves by mist net, with 69 bats resampled over the study for a total of 706 sampling instances. A total of 1,394 oral and rectal swabs were collected non-destructively and screened using molecular techniques for coronaviruses, paramyxoviruses, rhabdoviruses, flaviviruses, and filoviruses. Of these samples, 399 (56.5%) were collected during the rainy season and 307 (43.5%) during the dry season. Coronavirus RNA was detected in 59/706 (8.36%) of samples from *Rhinolophus spp.* (n = 35), *Hipposideros caffer* (n = 12), *Myonycteris angolensis* (n = 6), and *Miniopterus* spp. (n = 6). Six bats (0.85%) were positive for paramyxoviruses. Finally, (3 *H. caffer*, 1 *M. angolensis*, 1 *Rhinolophus spp.* and 1 *Nycteris thebaica*) 3 *Rhinolophus* bats were positive for rhabdoviruses (0.42%, all *Rhinolophus spp.*). No samples were positive for filovirus or flavivirus RNA. This project has generated novel data on the association of bat species and different viral strains present in these bats, advancing our knowledge of viral ecology and spillover risk at the human/bat interface.

## Introduction

Bats are ecologically and taxonomically diverse, with more than 1,500 species distributed globally (Turmelle and Olival 2009, Burgin 2018, Simmons & Cirranello. 2025). This makes them the second most diverse mammalian taxon to rodents, comprising about 20% of mammal species (Gupta et al. 2021). Bats contribute to ecosystem function by regulating insect populations, pollinating plants, and providing seed dispersal services, underscoring their socioeconomic importance to rural communities (Aggrey et al. 2024, Siya 2025). In Uganda, people have indicated support for bat conservation given the value of these ecological services and others provided by bats (Siya 2025). This ecological and socioeconomic value is challenged by the recognition of some bats as reservoirs for zoonotic diseases that pose significant threats to human and livestock health. These disease-causing pathogens include coronaviruses (Anthony et al. 2017), paramyxoviruses (Drexler et al. 2012), and filoviruses (Olival and Hayman 2014). Recognizing the importance of conserving bats and protecting people from bat-associated viruses, bat studies in Uganda and beyond have increasingly adopted a One Health framework, integrating surveillance, research, and intervention strategies to simultaneously protect human health, bat populations, and the surrounding ecosystem (Kayiwa 2024).

Most emerging and re-emerging infectious diseases are a result of interactions among humans and animals (Woolhouse and Gowtage-Sequeria 2005, Jones et al. 2008). The increasing human population and associated anthropogenic activities in Uganda have led to increased contact between people and wildlife, including bats, which have been implicated as reservoirs for several zoonotic pathogens in Uganda such as Marburg virus (family *Filoviridae*), Sosuga virus (family *Paramyxoviridae)*, and Kasokero virus (family *Nairoviridae*) (Kalunda et al. 1986, Towner et al. 2009, Amman et al. 2015). The interface of human-bat contact can threaten bat populations (Frick et al. 2020) and has created a need to understand the context and drivers behind the spillover of bat-borne infections into the human population. Pathogen transmission from bats to humans can occur via exposure to aerosols during cave trips i.e. histoplasmosis; Diaz 2018), ingestion of food sources contaminated by bats (Mickleburgh et al. 2009, Islam et al. 2016), or bites (i.e. rabies virus; Fenton et al. 2020). With the importance of protecting human and animal health, and global economies, infectious disease surveillance has been a key focus of national and international agencies for many years (Kading et al. 2018a). Recent large-scale outbreaks of emerging viruses have prompted research and surveillance activities to determine the importance of bats in harboring high-profile agents (Kupferschmidt 2018, Lau et al. 2020).

The drivers of disease emergence and biodiversity loss have a combined impact (Romanelli et al. 2015) warranting a coordinated approach to monitor, detect and mitigate changes in ecological conditions that may result in increased spillover risk. The linkage of biodiversity and infectious disease emergence has been recognized by the United Nations Convention of Biological Diversity and has led to efforts to support implementation of a Global Action Plan on biodiversity and health (CBD, 2025). Uganda is an emerging zoonotic disease hotspot (Jones et al. 2008, Nantima et al. 2019), particularly for bat-associated zoonoses (Towner et al. 2009, Amman et al. 2015, Ninsiima et al. 2024). This is particularly significant as Uganda is home to 98 bat species, with new species still being documented and contributing to its biodiversity (Monadjem et al. 2021); Simmons & Cirranello, 2025). Modelling studies indicate that Mount Elgon lies within one of the six hotspots expected to have a high diversity of bats (Herkt et al. 2016). It has been designated as a man and biosphere reserve, conveying the urgent need to balance human-environment interactions. The area also experiences land use and land cover changes with potential negative consequences for mammalian species like bats that are sensitive to such changes. In particular, deforestation, agricultural expansion and wood fuel demand are affecting the Mt Elgon ecosystem, although canopy cover is particularly important to preserving bat biodiversity (Bailey et al. 2019, von Kocemba et al. 2025). Thus, One Health surveillance initiatives are needed to protect bats and their habitats in Uganda and simultaneously protect human health from potential emerging virus spillover events.

Here, we describe a diversity of viruses detected in cave-dwelling bats across six sites in Eastern Uganda. Our data contribute information regarding viral detection amongst bat species sampled in both a temporal and spatial manner, given that we sampled during rainy and dry seasons. Viral phylogeny has also allowed us to assess genetic relatedness of other viruses previously detected in bats across multiple geographies. The outcomes of this research will be used to assess human and bat health risks, tailor conservation messaging, and inform One Health surveillance and response capacity needs at the national level.

## Methods

### Study area

Bats were captured for nonlethal sampling at six caves: five caves in Kapchorwa district, and one cave in Kween district, Uganda (Figure 1), from a study area described previously (Siya 2025). Cave names have been obscured using a coding system to protect the exact locations for conservation purposes. All caves were located outside the Mount Elgon National Park boundary, but within the Mount Elgon Transboundary Ecosystem. The average distance between the caves sampled in the study is 7.94 km, with a minimum distance of 1.15 km and a maximum of 16.8 km. This unique ecosystem comprises an expansive, extinct volcano spanning the border of Uganda and Kenya (UNESCO 2025) with many caves that provide critical habitat for bats and serve multiple purposes for local and tourist communities (Siya 2025).

**Figure 1.**
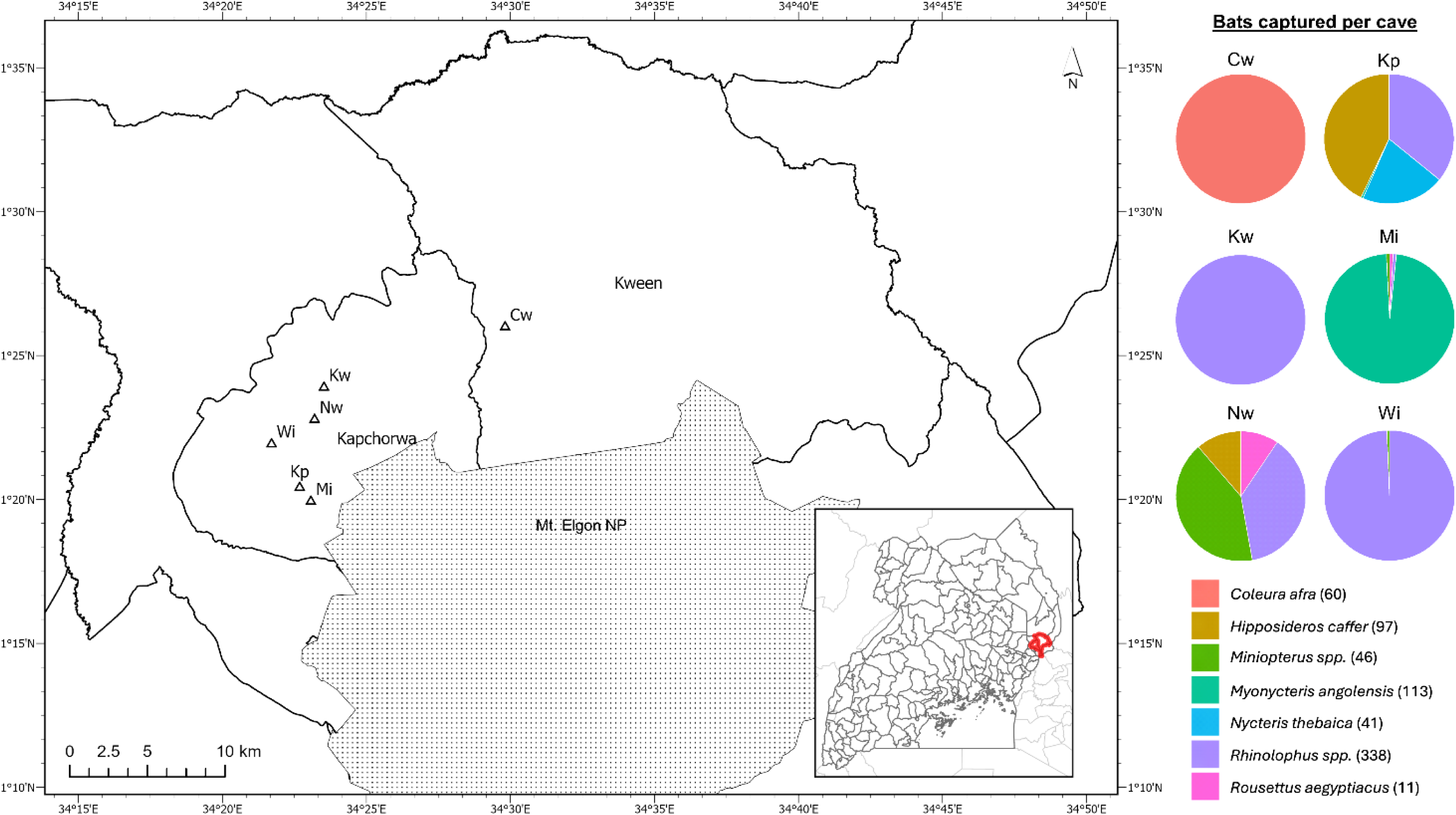
Bats were captured from six caves in the Kapchorwa and Kween districts of Uganda. Pie charts depict the proportion of species captured from each cave during the study. Note that captures were biased towards species of interest. Cave names are abbreviated to protect the locations of the caves for conservation purposes.

### Sample collection

Localities were sampled twice per year to capture any seasonal trends in viral shedding. Rainy season sampling occurred in May-June of 2021, 2022, and 2023. Dry season samples were collected in January of 2022 and 2023. Of the six caves, three caves were sampled for one wet and one dry season each, and the remaining three caves were sampled for three wet and two dry seasons over the course of the study. Bats were captured using mist nets or hand nets and placed in cloth holding bags until processing. Bat species from which we expected to find coronaviruses and other viral families of public health importance were selectively sampled. Bats were implanted with a Biomark Mini 8 (BIO8.B.04V1 PLT) Passive Integrated Transponder (PIT) tag (Biomark, Boise, ID, USA) prior to release at the cave entrance where they were captured. A 4mm wing punch was taken from a subset of bats and stored dry on silica gel desiccant for genetic species identification. Feces were also opportunistically collected from bats during sampling and stored on desiccant. Oral and rectal swab samples were collected via small FLOQ (oral) or MicroFLOQ (rectal) swabs (Copan Diagnostics, Murrieta, CA, USA). Duplicate oral and rectal swabs were stored in either Ambion MagMax Lysis Buffer (ThermoFisher Scientific, Waltham, MA, USA) or BD Universal viral transport media (VTM; Becton Dickinson, Franklin Lakes, NJ, USA) and placed directly onto dry ice, then removed to a liquid nitrogen dewar, and finally –80°C storage until sample processing at the Uganda Virus Research Institute (UVRI) in Entebbe, Uganda.

### Bat Species Identification

Bats were preliminarily identified based on morphological characteristics (i.e., forearm length, weight, and noseleaf width, in the case of horseshoe bats), following the identification key by Thorn & Peterhans (Thorn and Peterhans 2009). Certain genera (*Rhinolophus*, *Hipposideros*, *Myonycteris*, and *Nycteris*) contain one or more cryptic species complexes (Demos et al. 2019a, Demos et al. 2019b, Demos et al. 2020, Patterson et al. 2020), thus requiring additional species confirmation. To address taxonomic uncertainty for several genera, DNA was extracted from wing punches of representative bat species using DNeasy Blood & Tissue kits with an overnight proteinase K digestion (Qiagen; Venlo, Netherlands). For 4 *Nycteris* sp. bats and 1 *Hipposideros* bat from which no wing punch was collected, gDNA was extracted from silica desiccated feces usingQiagen’s QIAmp PowerFecal Pro DNA Kit. A total of 87 *Rhinolophus*, 19 *Hipposideros*, 4 *Nycteris*, and 20 *Miniopterus* spp. samples were subjected to PCR amplification of a portion of the mitochondrial cytochrome *b* gene (*cytb*) for further molecular identification of captured species following pre-established protocols (Bickham et al. 2004, Demos et al. 2019a). PCR product was purified using NEB Monarch PCR & DNA Cleanup kit prior to Sanger sequencing (NEB; Ipswitch, MA and Azenta; Burlington, MA). Sanger sequencing output reads were trimmed and assembled into contigs using Geneious Prime (ver. 2019.2.1)(Boston, MA). Consensus sequences were aligned using the MAFFT plugin (ver 1.5.0) in Geneious Prime alongside representative reference sequences for each genus (Demos et al. 2019a, Demos et al. 2019b, Demos et al. 2020, Patterson et al. 2020). Maximum likelihood estimates of *cytb* gene trees were generated using IQ-TREE (ver 2.4.0)(Trifinopoulos et al. 2016), including a built-in tool to identify the optimal maximum likelihood substitution model. We performed an ultrafast branch support analysis with 1000 bootstraps and maximum 1000 iterations, and final trees were visualized in iTOL (ver 7.5) (Letunic and Bork 2024).

### RNA extraction and PCR methods

RNA was extracted from the rectal (n = 693) and oral (n = 701) swabs using the MagMAX Viral/Pathogen Nucleic Acid Isolation Kit (ThermoFisher Scientific, Waltham, MA, USA) according to manufacturer’s instructions. RNA was reverse transcribed to cDNA using the SuperScript IV First-Strand Synthesis System (Invitrogen,Carlsbad, CA). An internal control targeting the cytochrome *b* gene was used for detecting amplifiable nucleic acid material in the RNA extracts (Ngo and Kramer 2003). The cDNA was then subjected to PCR using consensus primers targeting the NS5 gene of flaviviruses (Moureau et al. 2007), the pol (L) gene of paramyxoviruses (Tong et al. 2008), and the RNA-dependent RNA polymerase (Rdrp) gene of coronaviruses (Watanabe et al. 2010, Quan et al. 2010), the L gene of filoviruses (Zhai et al.

2007), and the L gene of rhabdoviruses (Balinandi et al. 2022). The PCR products were visualized under UV light after gel electrophoresis. Those bands with intensity correlating to transcript abundance and expected target size were confirmed by Sanger or Next-Generation Sequencing (NGS) amplicon sequencing.

### Sequencing and analysis of presumptively positive RNA samples

#### Sanger Sequencing

Putatively positive PCR amplicons from all assays representing sampling trips from May 2021 through January 2023 were transported to Colorado State University. Amplicons were purified using the Monarch PCR & DNA Cleanup kit (NEB; Ipswitch, MA). Products were then subjected to A-tailing using a generalized protocol wherein amplicons were incubated in a solution of 10X Taq Polymerase Buffer (Invitrogen; Waltham, MA), 0.5U of Taq Polymerase (Invitrogen; Waltham, MA), 1mM dATP (Zymo; Orange, CA), and PCR-grade water (Ambion; Waltham, MA) to a final volume of 50µL for 20 minutes at 72°C. Amplicons were then inserted into a pCR4 sequencing vector using TOPO TA cloning methods as specified by the manufacturer (Invitrogen; Waltham, MA). Transformants were screened by PCR using plasmid-based M13F and M13R priming sites. Up to 5 positive transformants were selected for growth in LB broth under antibiotic selection, followed by plasmid extraction using a Zymo ZR Plasmid Miniprep kit (Zymo; Orange, CA). Extracted samples were submitted to Azenta (Burlington, MA) for Sanger sequencing. All sequenced products were analyzed using Geneious Prime (Boston, MA) by identifying and removing plasmid and amplicon-specific primer sequence. Sequences were then subjected to BLAST (Altschul et al. 1990, Altschul et al. 1997, Zhang et al. 2000, Morgulis et al. 2008, Camacho et al. 2009) to determine positive or negative representation of viral sequences.

#### Next-Generation Sequencing

All presumptive PCR positive amplicons obtained from bats sampled during the May 2023 trip were amplicon-sequenced and analyzed at the UVRI genomics laboratory. PCR amplicons were subjected to a 1:1 ratio bead-clean-up using AMPure XP magnetic beads (Beckman Coulter, Brea, CA, USA). Amplicon concentration was determined using the Qubit™ dsDNA High Sensitivity Assay (Thermo Fisher Scientific, Waltham, MA, USA) on a Qubit™ 4.0 fluorometer following manufacturer’s recommendation.

Paired-end libraries were prepared using the TruSeq® Stranded Total RNA Library Prep (Illumina, Inc. San Diego, CA, USA), starting at the A-tailing step, and proceeding with other downstream steps as recommended by the manufacturer. The libraries were then quantified using the Qubit™ dsDNA High Sensitivity Assay and fragment analysis performed using the Agilent High Sensitivity D1000 ScreenTape assay (Agilent Technologies, Santa Clara, CA, USA). The libraries were normalized and pooled, then denatured using 0.2N NaOH diluted using pre-chilled HT1 buffer to make a loading concentration of 12pM on the Illumina Miseq (2 x 250 cycles).

The generated sequence data were quality trimmed (Q30) to remove adapters and low-quality reads using trim galore v0.6.10 (Krueger et al. 2023) and trimming results assessed and visually inspected using FastQC v0.12.1 (Ewels et al. 2016) and multiqc v1.14 (Ewels et al. 2016). Assembly of the reads was then performed using SPAdes v3.15.5 and the taxon of the longest and highest depth contig determined using BLAST v2.16.0+ (Prjibelski et al. 2020) with the NCBI core nucleotide database. All sequences are available in GenBank (Supplementary Table 1).

### Phylogenetic analysis

The sequences from NGS were compiled by viral family together with those from Sanger sequencing and analyzed phylogenetically to investigate diversity among families. Multiple Sequence Alignment (MSA) was performed for sequences of each of the viral families using MAFFT v7.526 (Camacho et al. 2009) (1000 iterations). Closely related sequences from GenBank and RefSeq were added to the alignment for reference. Shorter sequences were aligned separately from longer sequences and their alignment added onto the longer backbone using the option “--add fragments” to avoid false gap opening. Maximum likelihood phylogenetic trees were then constructed from the alignments using IQ-TREE v3.0.1 (Minh et al. 2020) (1000 bootstrap alignments) with the best fit substitution model chosen based on the Bayesian Information Criterion (BIC). The trees were then edited in iTOL v7 (Letunic and Bork 2024).

### Recombination analysis for coronaviruses

Potential recombination events were evaluated in Geneious Prime® (ver 2025.0.3). Each query sequence was aligned against its most closely related reference sequence from the NCBI RefSeq database using pairwise global alignments with default parameters. The resulting alignments were inspected visually to assess local sequence identity across the sequences. Regions of markedly reduced pairwise identity were noted as potential recombination breakpoints. To confirm these, sequences were divided at the inferred breakpoint positions, and each fragment was realigned to its closest RefSeq reference sequence. Better pairwise alignments of the fragments with different references or regions of the same reference were taken as supporting evidence of recombination.

## Results

### Bat captures

635 unique individual bats were captured and tested during the study, which with recaptures across five field seasons, resulted in 706 captures. Bats were captured from a total of six caves across the Mt. Elgon region of Eastern Uganda. Bats captured represented 7 genera: 1) *Coleura afra* (n = 60), 2) *Hipposideros caffer* (n = 94), 3) *Miniopterus* spp. (n = 46), 4) *Myonycteris angolensis* (n = 63), 5) *Nycteris thebaica* (n = 40), 6) *Rhinolophus* spp. (n = 321), and 7) *Rousettus aegyptiacus* (n = 11). Approximately 52.49% (369) of total bats sampled were female, while 47.51% (334) were male, suggesting even sample representation across sexes (Table 1). Age determination of captured bats showed that approximately 80.2% were classified as adults, while 17.2% were sub-adult, and 2.6% juveniles. 56.52% (399) of captures occurred in the rainy season (May) and 43.48% (307) during the dry season (January). However, not all caves were sampled on each collection trip; caves were removed or added to the study in collaboration with people local to the region. Cave Cw was sampled only in May 2021 and January 2022. Caves Kw and Nw were sampled in January 2023 and May 2023. Caves Kp, Mi, and Wi were sampled across all 5 collection trips. Thus, some caves were sampled twice while others were sampled five times, which impacts interpretation of recapture rates from each cave.

**Table 1.**
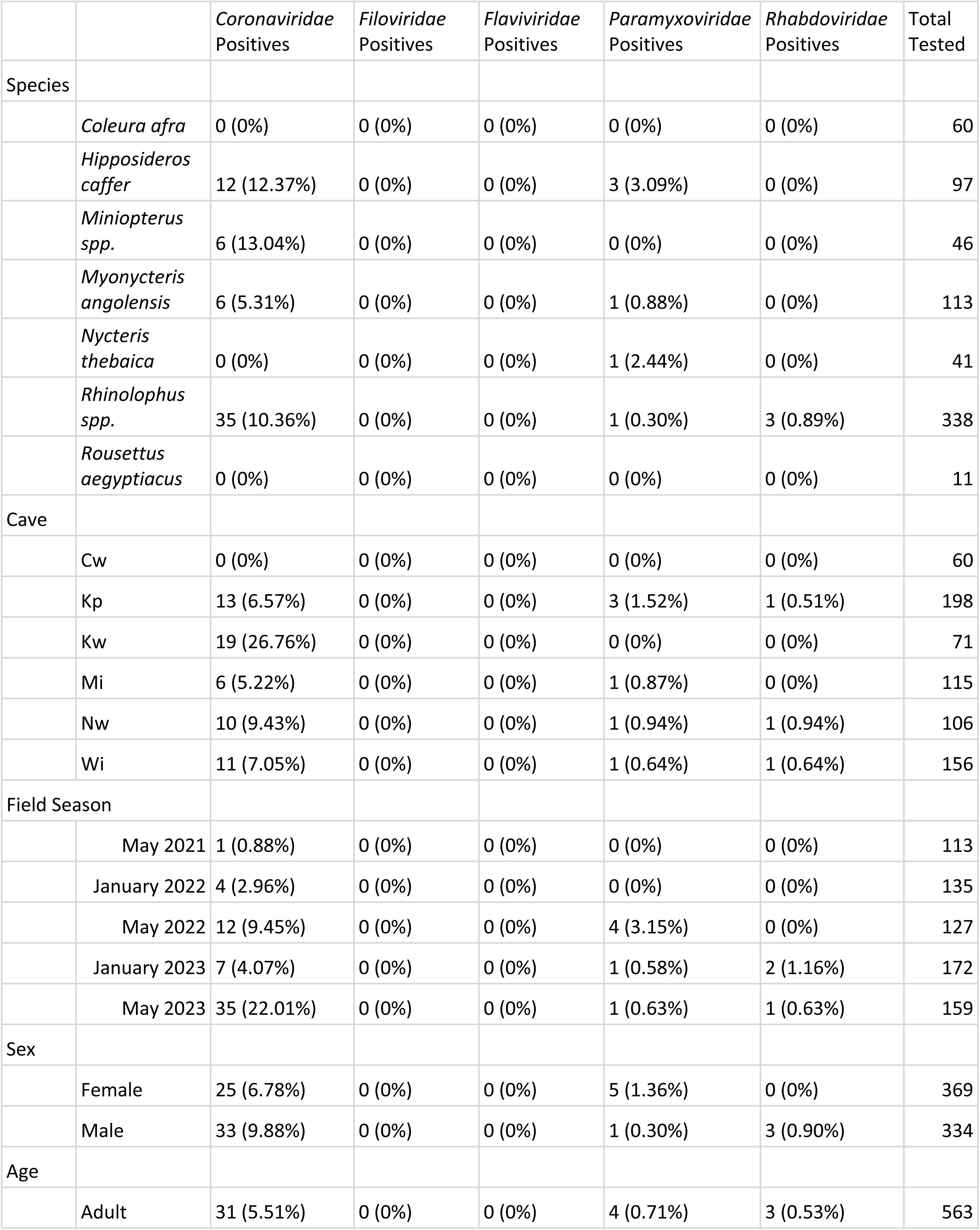
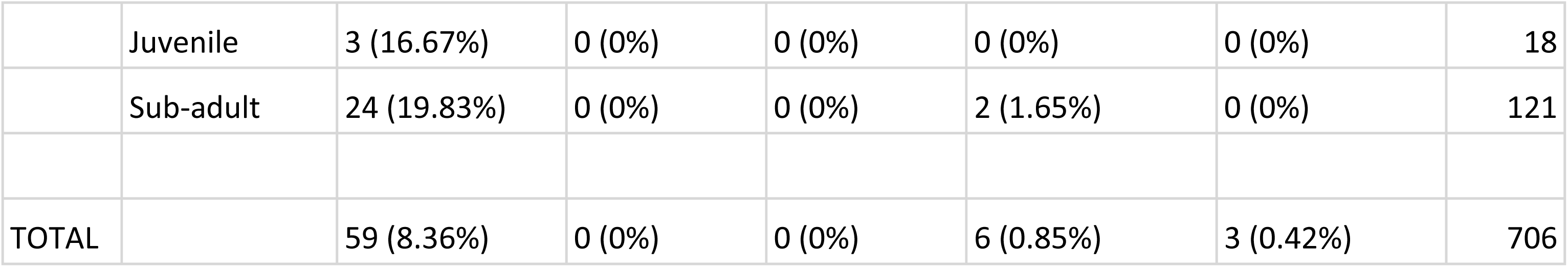
Family-level virus detections based on PCR assays and confirmed by sequencing. The raw number of positive captures followed by the percentage of positives based on the total number of captures are presented here.

|  |  | Coronaviridae<br>Positives | Filoviridae<br>Positives | Flaviviridae<br>Positives | Paramyxoviridae<br>Positives | Rhabdoviridae<br>Positives | Total<br>Tested |
| --- | --- | --- | --- | --- | --- | --- | --- |
| Species |  |  |  |  |  |  |  |
|  | *Coleura afra* | 0 (0%) | 0 (0%) | 0 (0%) | 0 (0%) | 0 (0%) | 60 |
|  | *Hipposideros caffer* | 12 (12.37%) | 0 (0%) | 0 (0%) | 3 (3.09%) | 0 (0%) | 97 |
|  | *Miniopterus spp.* | 6 (13.04%) | 0 (0%) | 0 (0%) | 0 (0%) | 0 (0%) | 46 |
|  | *Myonycteris angolensis* | 6 (5.31%) | 0 (0%) | 0 (0%) | 1 (0.88%) | 0 (0%) | 113 |
|  | *Nycteris thebaica* | 0 (0%) | 0 (0%) | 0 (0%) | 1 (2.44%) | 0 (0%) | 41 |
|  | *Rhinolophus spp.* | 35 (10.36%) | 0 (0%) | 0 (0%) | 1 (0.30%) | 3 (0.89%) | 338 |
|  | *Rousettus aegyptiacus* | 0 (0%) | 0 (0%) | 0 (0%) | 0 (0%) | 0 (0%) | 11 |
| Cave |  |  |  |  |  |  |  |
|  | Cw | 0 (0%) | 0 (0%) | 0 (0%) | 0 (0%) | 0 (0%) | 60 |
|  | Kp | 13 (6.57%) | 0 (0%) | 0 (0%) | 3 (1.52%) | 1 (0.51%) | 198 |
|  | Kw | 19 (26.76%) | 0 (0%) | 0 (0%) | 0 (0%) | 0 (0%) | 71 |
|  | Mi | 6 (5.22%) | 0 (0%) | 0 (0%) | 1 (0.87%) | 0 (0%) | 115 |
|  | Nw | 10 (9.43%) | 0 (0%) | 0 (0%) | 1 (0.94%) | 1 (0.94%) | 106 |
|  | Wi | 11 (7.05%) | 0 (0%) | 0 (0%) | 1 (0.64%) | 1 (0.64%) | 156 |
| Field Season |  |  |  |  |  |  |  |
|  | May 2021 | 1 (0.88%) | 0 (0%) | 0 (0%) | 0 (0%) | 0 (0%) | 113 |
|  | January 2022 | 4 (2.96%) | 0 (0%) | 0 (0%) | 0 (0%) | 0 (0%) | 135 |
|  | May 2022 | 12 (9.45%) | 0 (0%) | 0 (0%) | 4 (3.15%) | 0 (0%) | 127 |
|  | January 2023 | 7 (4.07%) | 0 (0%) | 0 (0%) | 1 (0.58%) | 2 (1.16%) | 172 |
|  | May 2023 | 35 (22.01%) | 0 (0%) | 0 (0%) | 1 (0.63%) | 1 (0.63%) | 159 |
| Sex |  |  |  |  |  |  |  |
|  | Female | 25 (6.78%) | 0 (0%) | 0 (0%) | 5 (1.36%) | 0 (0%) | 369 |
|  | Male | 33 (9.88%) | 0 (0%) | 0 (0%) | 1 (0.30%) | 3 (0.90%) | 334 |
| Age |  |  |  |  |  |  |  |
|  | Adult | 31 (5.51%) | 0 (0%) | 0 (0%) | 4 (0.71%) | 3 (0.53%) | 563 |
|  | Juvenile | 3 (16.67%) | 0 (0%) | 0 (0%) | 0 (0%) | 0 (0%) | 18 |
|  | Sub-adult | 24 (19.83%) | 0 (0%) | 0 (0%) | 2 (1.65%) | 0 (0%) | 121 |
| TOTAL |  | 59 (8.36%) | 0 (0%) | 0 (0%) | 6 (0.85%) | 3 (0.42%) | 706 |

### Molecular species confirmation of bats

Given the cryptic nature of certain species collected in this study, molecular identification was completed by sequencing *cytb* from DNA extracted from wing biopsy samples. *Rhinolophus* spp. bats all fell within clade 1 of the *Rhinolophus fumigatus / eloquens* cryptic species complex (Demos et al. 2019a). The species of *Hipposideros* bats observed were identified as belonging to *H.caffer* species complex lineage 8. *Miniopterus* spp. bats morphologically resembled *M. fraterculus* but *cytb* sequences fell into 3 distinct clades and are thus referred to as *Miniopterus* spp. because genomic confirmation was not possible for all individuals. (Supplementary Fig 1). *Nycteris thebaica* bats were confirmed as belonging to lineage 3 within the species complex. All *cytb* sequences generated in this study are available in GenBank accession numbers PX734756 – PX734885 (Supplemental Table 2).

### Bat recaptures

There were 69 recapture events of PIT-tagged bats over the course of the study. These 69 recaptures comprised 49 individual bats: 31 bats were recaptured once, 7 bats were captured three times each, and 11 bats (all *Myonycteris*) were captured four times each. Within caves Cw, Kw, and Nw, there were no bats recaptured from the same cave in which they were originally marked. The recapture rates of the remaining caves were as follows: Kp n=14 (10.85%), Mi n=50 (43.86%), Wi n=5 (3.55%). Across all six caves, the average recapture rate was 11.26%. *M. angolensis* individuals represented a smaller colony in cave Mi and were recaptured more frequently. Assessment of cross-cave recaptures showed that five bats, all male *Rhinolophus* spp. bats, were recaptured at a different cave from where they were first captured. Four of these bats were recaptured in different seasons (though always within the same year), while one captured in Kw was recaptured in Nw the following day. These recaptures demonstrate connection between Wi, Kp, Kw, and Nw caves all four major *Rhinolophus-*inhabited caves sampled in the study. One of these recaptures was from a bat originally PIT tagged at a cave not sampled in this study, showing connection with at least one additional cave beyond this study. One recaptured bats tested positive for a coronavirus during the May 2023 capture at Nw. Additionally, there is evidence of one adult female *M. angolensis* bat switching from cave Mi to an unknown cave based on GPS tracking data (data not shown). This individual also tested positive for a coronavirus and is the only bat to provide direct evidence of cave swapping with detectable coronavirus.

### Viral RNA detections

All bat oral and rectal swabs were screened for viruses in the families *Coronaviridae, Filoviridae*, *Orthoflaviviridae*, *Paramyxoviridae, and Rhabdoviridae* (Table 1, Figure 2). No RNA resulting from swabs were positive for filoviruses or flaviviruses. 68 individual bats were positive for viral RNA in rectal and/or oral swabs for coronaviruses (n = 59), paramyxoviruses (n = 6), and rhabdoviruses (n = 3) (Figure 2). No individual bats were positive for more than one viral family, and no recaptured bats were positive longitudinally across the sampling period. Of the viral detections made, 13.5% (n = 54/399) were during the rainy season and 4.6% (n = 14/307) during dry season sampling.

**Figure 2.**
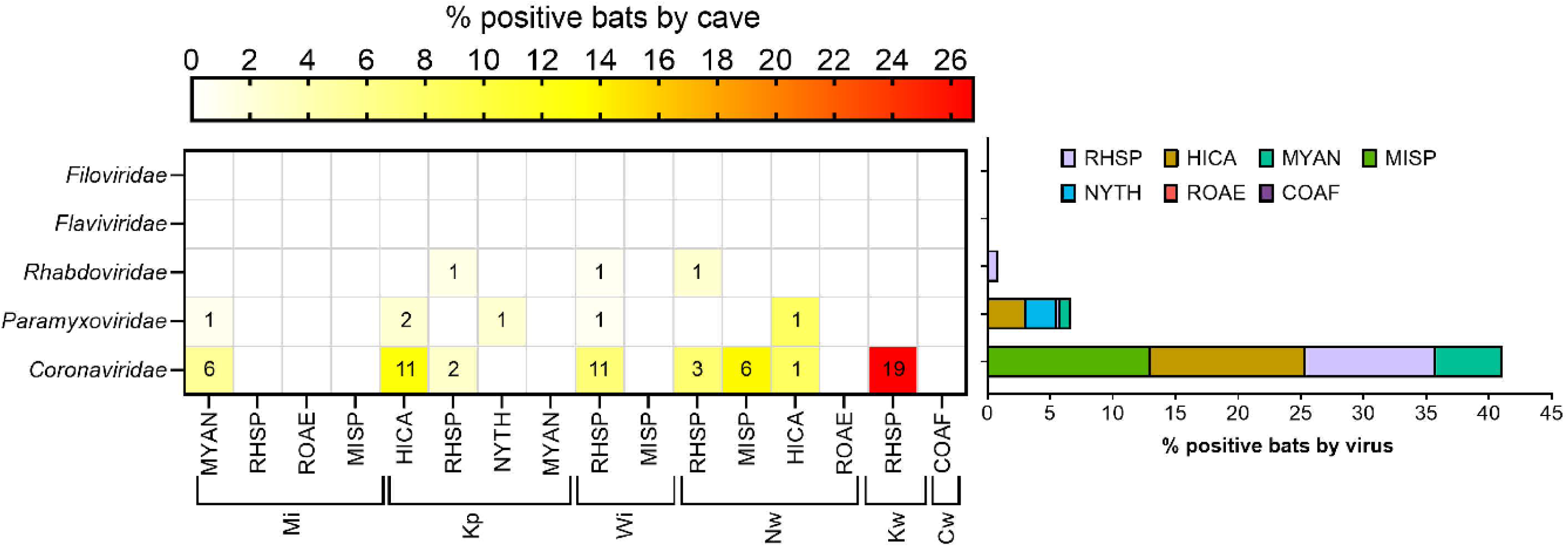
Virus families detected from different bat species and caves. The numbers in the heat map represent the number of individual bats that tested positive for each species and cave. Cave designations are as in Figure 1. RHSP = *Rhinolophus* spp.; MISP = *Miniopterus* spp.; HICA = *Hipposideros caffer*; NYTH = *Nycteris thebaica*; MYAN = *Myonycteris angolensis*; ROAE = *Rosettus aegyptiacus*; COAF = *Coleura afra*.

### Coronaviruses

Coronaviruses were detected across five of the six caves for at least one sampling trip, revealing an 8.36% (59/706) overall detection of viral RNA among bats included in this study (Figures 3 and 4). The highest percentage of infected bats was observed in *Miniopterus* spp. (n = 6/46; 13.04%), all of which were collected from the same cave during a single dry season. Phylogenetic reconstruction of two portions of the ORF1 from coronaviruses revealed the RNA was similar among positive bats and belonged to the genus *Alphacoronavirus* and subgenus *Minunacovirus*. Sequences clustered closely with other minunacoviruses detected in other *Miniopterus* spp. bats in Kenya and China.

**Figure 3.**
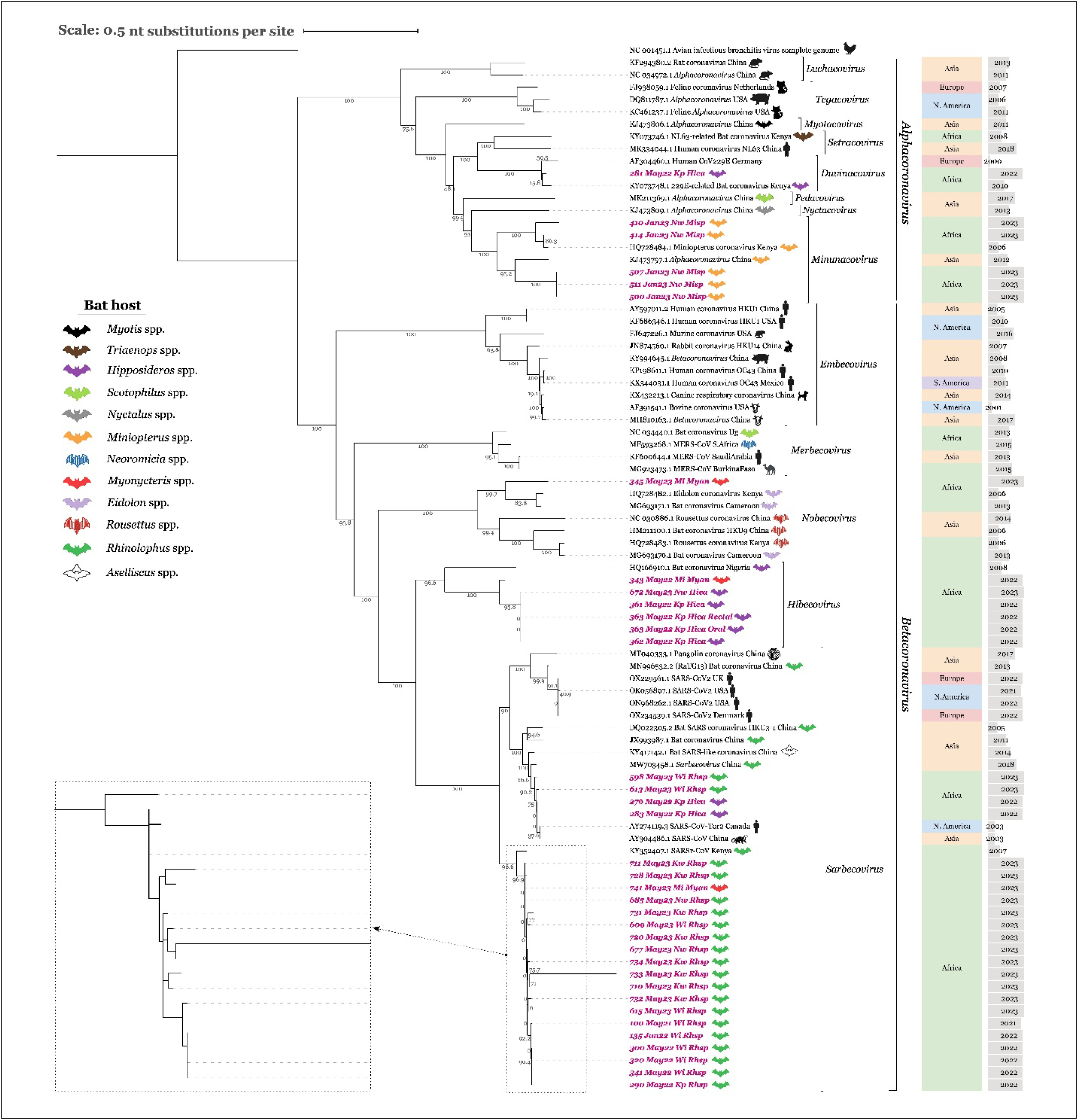
Maximum likelihood tree based on nucleotide alignment of coronaviruses amplicons detected across study caves from *Rhinolophus, Hipposideros, Myonycteris*, and *Miniopterus* bats. Aligned sequences represent a conserved 400-nt fragment of the Rdrp gene (Quan, 2010). The tree is rooted by Avian infectious bronchitis virus. Sequences detected in the current study are highlighted with pink text. Branch support values represent bootstrap percentages based on 1,000 replicates. All accession numbers stated are listed in the NCBI nucleotide database. Icons right of the branch labels represent host species of the virus, along with the continent where the sample was collected and the collection year.

**Figure 4.**
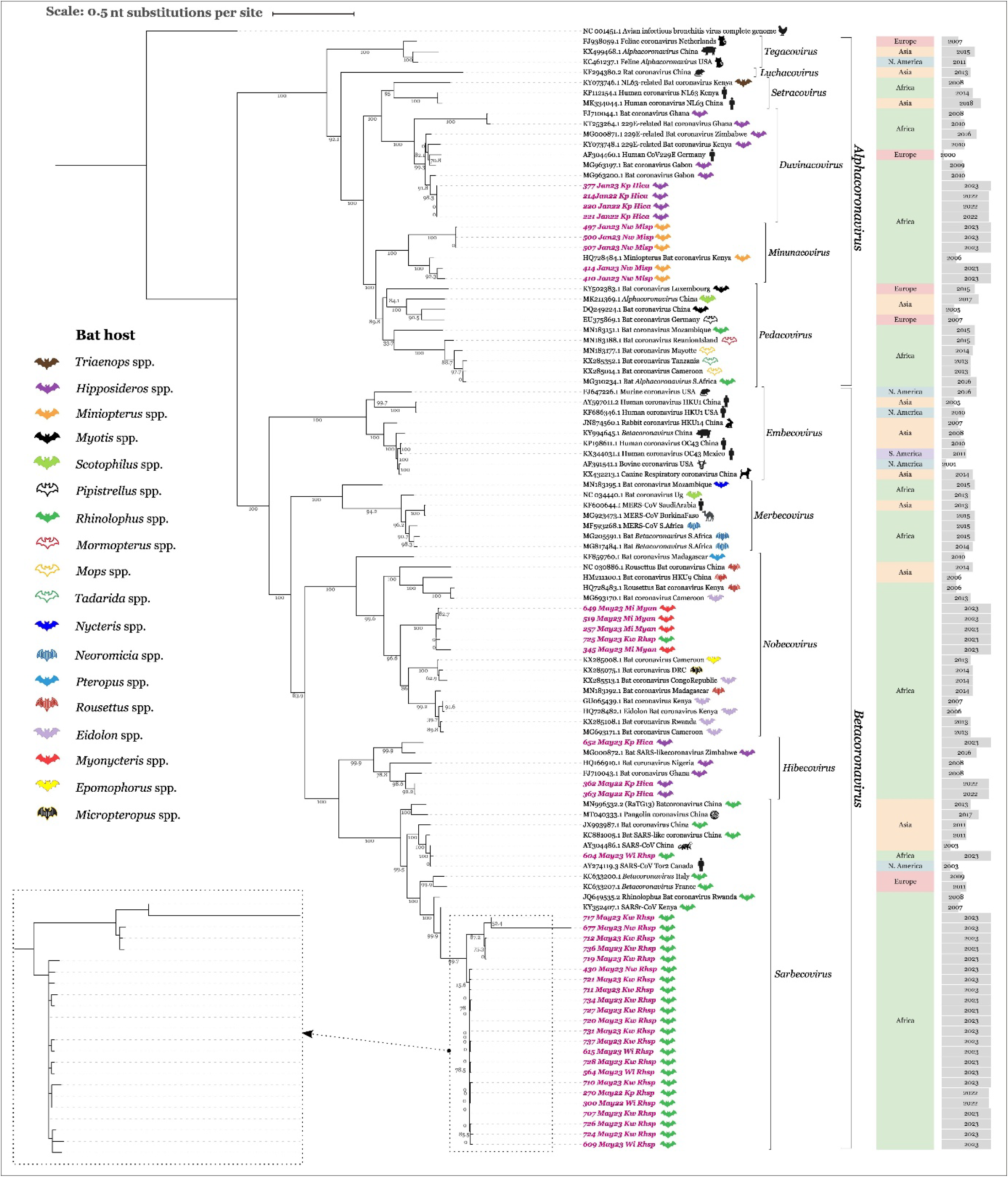
Maximum likelihood tree based on nucleotide alignment of coronavirus amplicons detected across study caves from *Rhinolophus, Hipposideros, Myonycteris*, and *Miniopterus* bats. Aligned sequences represent a conserved 440-nt fragment of the Rdrp gene (Watanabe, 2010). The tree is rooted by Avian infectious bronchitis virus. Sequences detected in the current study are highlighted with pink text. Branch support values represent bootstrap percentages based on 1,000 replicates. All accession numbers stated are listed in the NCBI nucleotide database. Icons right of the branch labels represent host species of the virus, together with the continent on which the sample was collected and the collection year.

*Hipposideros caffer*, which were sampled across two study caves, represented the species with the second highest positivity rate for coronaviruses, at 12.37% (12/97). Eleven out of the twelve positive bats were sampled in Cave Kp, with the other singular detection originating from Cave Nw. Coronaviruses were detected in *H. caffer* across all sampling time points, except May 2021. Five of the coronavirus positive bats from cave Kp had high sequence similarity to each other, even though detections were made across multiple seasons (i.e., January 2022, May 2022, and May 2023). Phylogenetically, these sequences were similar to other 229E-like alphacronaviruses within the subgenus *Duvinacovirus*. Of the additional seven bats with positive RNA for coronaviruses, all were similar to other betacoronviruses. Five of these bats produced viral sequences that grouped within the subgenus *Hibecovirus* and were observed across two dry seasons (May 2022 versus May 2023) in two distinct caves. Lastly, two instances of viral RNA detections from *H. caffer* in May 2022 produced sequences highly similar to various other bat-associated sarbecoviruses and also similar to detections from *Rhinolophus* spp. within this study.

*Rhinolophus* spp. bats represented 10.36% (35/338) of the total coronavirus detections made in this study and were detected longitudinally across all sampling time points (May 2021-May 2023). Coronavirus detections were made from four of the five caves where members of the *Rhinolophus* genus were observed. All RNA-detections from *Rhinolophus* spp. bats clustered within the genus *Betacoronavirus.* Thirty-four of the positive bats were highly similar to each other and to other *Rhinolophus* spp.-associated *Sarbecovirus*. Only one sequence from a *Rhinolophus* sp. bat sampled in May 2023 in Cave Kw was similar to other sequences within the subgenus *Nobecovirus*. Other detections within *Nobecovirus* came from *M. angolensis* sampled in this study.

All *M. angolensis* were observed in Cave Mi and 5.31% (6/113) of samples tested were positive for coronavirus RNA. All sequence data suggested the coronaviruses detected were betacoronaviruses. Four of the six sequences clustered most closely within the subgenus *Nobecovirus* and were like other bat-associated nobecoviruses from across Africa. Additional *M. angolensis* bats were positive for coronavirus RNA most closely resembling viruses within the *Hibecovirus* and *Sarbecovirus* subgenera. The latter detection of a *M. angolensis*-associated *Sarbecovirus* shows high similarity with other sarbecoviruses detected in *Rhinolophus* spp. that were collected in May 2023.

Nineteen (26.76%) of coronavirus positives came from cave Kw, all from May 2023. While this was a high detection rate for one season, cave Kw was only sampled later in the study period so there are more limited dates from this location for comparison. Additionally, the frequency of positives was not evenly distributed across age classification. Forty-six percent of coronavirus positives were from juvenile or sub-adult bats, which made up only 20% of bats tested. Rectal swabs were more often positive for coronavirus RNA than oral swabs; 6 oral swabs were positive for coronavirus RNA and 54 rectal swabs were positive for coronavirus RNA. Additionally, screening samples for coronavirus RNA using two complementary consensus assays increased coronavirus detection rates. If only one of these assays had been used, many positive samples would have been missed. Thirty-five bats were positive in one assay (Quan et al. 2010): 4 from oral swabs, 34 from rectal swabs, and 1 bat positive in both oral and rectal swabs. A total of 41 bats were positive by the second assay (Watanabe PCR assay): 1 from an oral swab and 40 from rectal swabs. Seventeen rectal swabs were positive by both assays. No swabs were positive across all four-sample type and assay combinations.

### Recombination events among coronaviruses

The phylogenetic analysis exhibited exceptionally long branches in two betacoronavirus sequences, bat 733 (Figure 3) and bat 677 (Figure 4), which, when further investigated, revealed homologous recombination events. Using the reference genome KY352407.1, we characterized those potential recombination events in each strain.

The sequence 733-May23-Kw-Rhsp (209 bp) was identified as a potential recombinant, with a major parent contributing 131 bp (85.3% identity) and a minor parent contributing 77 bp (77.5% identity) in a reverse-complement orientation. The identified breakpoints were at positions 18588-18716 bp and 18449-18528 bp, respectively. The sequence 677-May23-Nw-Rhsp (398 bp) was also identified as a potential recombinant, with a major parent contributing 268 bp (91.2% identity) and a minor parent contributing 129 bp (92.9% identity). The breakpoints for this event were at positions 15363-15631 bp and 15528-15657 bp, respectively.

### Paramyxoviruses

A total of six bat rectal swab samples were positive for paramyxovirus RNA. These samples were collected across four of the six study caves. Three of positive samples were taken from *H. caffer* inhabiting Caves Kp and Nw and represented different sampling time points, showing longitudinal viral circulation of paramyxovirus strains. All viral sequences from *H. caffer* clustered with other parajeilongviruses detected in *Hipposideros* spp. in Africa and China. Additionally, there was a singular detection in *N. thebaica* sampled in May 2022 that demonstrated similarity to other parajeilongviruses. This detection was also made in Cave Kp, further highlighting circulation of multiple strains of parajeilongviruses among different bat species inhabiting the same cave. A bat from the genus *Rhinolophus* sampled in Cave Wi produced a positive RNA sequence that most closely resembled an orthorubulavirus, of which phylogenetic reconstruction showed to be similar to South American swine and bats. We also obtained a paramyxovirus positive sequence from a *M. angolensis* captured in Cave Mi in May 2022. Sequence analysis revealed high similarity to the genus *Henipavirus* and other such samples collected from *Rhinolophus* and *Eidolon* spp. from Australia and Africa, respectively. All paramyxovirus RNA detections were from rectal swabs; urine was not collected in this study.

### Rhabdoviruses

Sequencing and alignment of a portion of the rhabdovirus RdRp revealed three positive RNA samples across three different caves. All positive samples were collected from *Rhinolophus* spp., two in January 2023 and one in May 2023. Two of the three rhabdovirus-positive detections came from oral swabs, and one was from a rectal swab. Phylogenetic analysis showed that a *Rhinolophus* spp. sample collected in Cave Nw was closely related to members of the genus *Vesiculovirus*. Another *Rhinolophus* spp. sample collected in Cave Kp mostly closely matches other sequences from bats and tapeworms within the genus *Betaplatrhavirus*. A third bat captured in Cave Wi seemingly represents a unique virus with an unclear genus association.

## Discussion

Caves in the Mount Elgon region experience a high degree of human disturbance from the local human population as well as tourists, providing a dynamic interface for potential exposure of people to viruses carried by bats. Moreover, infectious disease surveillance capacity is considerably lacking in this geographical area (Siya et al. 2021). With the notable exception of Marburg virus in Egyptian rousette bats (*Rousettus aegyptiacus*) (Towner et al. 2007, Towner et al. 2009, Letko et al. 2020), little is known about the diversity and ecology of viruses circulating among cave-dwelling bats in Uganda and their potential public health impacts. Several bat taxa commonly found roosting in caves in this region are associated with viruses of known or potential public health significance, making it an ideal location to assess spillover risk: *Rhinolophus* (coronaviruses; Anthony et al. 2017, Latinne et al. 2024); *Hipposideros* (coronaviruses; Meyer et al. 2024); *Rousettus* (filoviruses, paramyxoviruses; (Amman et al. 2015, Markotter et al. 2019, Amman et al. 2024), and *Miniopterus* (filovirus; Kemenesi et al. 2018). Therefore, we sought to describe the diversity of viruses that people may be exposed to in a cave environment as a first step towards understanding potential health threats in this region.

### Viral detections

Multiple unique strains of alpha– and betacoronaviruses, paramyxoviruses, and rhabdoviruses were detected over a 2.5 year period. Viruses in the families *Coronaviridae, Paramyxoviridae*, and *Rhabdoviridae* are three of the most common virus families reported from bats (Letko et al. 2020). In this study, bats in the genera *Rhinolophus*, *Myonycteris*, and *Hipposideros* were each associated with viruses from two or three of these viral families (Figure 2 and Table 1), although no individual bats were found co-infected with viruses in these families. Oral and rectal swab samples collected from bats were also screened for filoviruses and flaviviruses, but no positive results were found for either of these viral families. The lack of flavivirus detections was somewhat surprising, since several no-known-vector flaviviruses are associated with bats in Africa, including Entebbe bat virus which was first isolated from the salivary glands of bats in Uganda (Lumsden et al. 1961, Kading et al. 2015, Blitvich and Firth 2017). Additionally, a high antibody prevalence to flaviviruses was previously reported from bats in Uganda, including those captured in this area (Kading et al. 2018b). We only tested a limited number of Egyptian rousette bats, which are known to harbor Marburg (*Filoviridae*) and Sosuga (*Paramyxoviridae*) viruses in Uganda (Towner et al. 2009, Amman et al. 2015).

### Coronaviruses

Alpha and betacoronaviruses detected in this study were similar to previously published strains detected in Africa. Alphacoronaviruses associated with Sundevall’s roundleaf bats (*Hipposideros caffer)* clustered in the *Duvinacovirus* subgenus (Figs 3-4). The genus *Duvinacovirus* contains human coronavirus strain 229E (van der Hoek et al. 2004), which is believed to have originated from hipposiderosid bats (Corman et al. 2015). Virus strains like 229E have been previously detected in *H. caffer, Rhinolophus*, and *Nycteris* spp. in Zimbabwe (Bourgarel et al. 2018, Chidoti et al. 2022); *Hipposideros* spp. in Cameroon and Guinea (Meta Djomsi et al. 2023); *H. ruber* in the Republic of Congo (Kumakamba et al. 2021); *Hipposideros ruber* in Gabon (Maganga et al. 2020), and *Hipposideros ruber* in Cameroon (Ntumvi et al. 2022). Our results are consistent with these reports from across Africa and provide additional data points representative of Uganda.

The high prevalence of coronaviruses in bent-winged bats (genus *Miniopterus;* Figure 2) was surprising given our sample size, however, there are many other reports of diverse coronaviruses in *Miniopterus* spp. bats. The coronavirus strains we detected in *Miniopterus* cluster phylogenetically in subgenus *Minunacovirus*, in the genus *Alphacoronavirus*. These strains are very similar to *Miniopterus* bat coronaviruses detected in Kenya (Tong et al. 2009), Zimbabwe (Chidoti et al. 2022), Cameroon (Meta Djomsi et al. 2023), Republic of Congo (Kumakamba et al. 2021), and also HKU7 and HKU8 coronaviruses described from *Miniopterus* spp. bats in Hong Kong (Poon et al. 2005). Chidoti et al. 2022 also reported alphacoronaviruses from *Miniopterus* that cluster in the *Decacovirus* subgenus, and the *Setravirus* subgenus with human CoV strain NL63. These virus strains from bent-winged bats are in different subgenera than human coronavirus strains NL63 and 229E (Fig 3); the human pathogenic potential of the minunacovirus strains detected in this study is unknown.

Sarbecoviruses (genus *Betacoronavirus*) were detected in bats in the genera *Rhinolophus*, *Hipposideros*, and *Myonycteris*. This finding is consistent with the global recognition that *Hipposideros* and *Rhinolophus* are associated with SARS-related coronaviruses (SARSr-CoVs) (Anthony et al. 2017, Wang et al. 2025). SARS-related betacoronaviruses have also been detected in these genera of bats from across Africa. A SARSr-CoV was detected in *Hipposideros caffer* in Zimbabwe (Bourgarel et al. 2018); Bourgarel et al. (2018) also reported circulation of alphacoronaviruses and a jeilongvirus-related (Paramyxovirus) among *Hipposideros* bats.

One of these cave sites from Zimbabwe and a different site were visited later by Chidoti et al., 2022 for further study of coronavirus ecology. Multiple strains of coronaviruses spanning alpha– and betacoronaviruses were detected, including multiple strains of sarbecoviruses from *Rhinolophus* spp. bats. Hibecoviruses were found in *Hipposideros, Rhinolophus*, and *Nycteris* spp. Merbecoviruses were detected in *Hipposideros, Nycteris,* and *Rhinolophus* co-roosting in the Magweto cave site. Meta Djomsi et al., 2023 detected sarbecoviruses in *Rhinolophus* from Cameroon and Guinea and hibecoviruses in *Hipposideros* (Meta Djomsi et al. 2023). Nziza et al., 2020 reported a sarbecovirus in *Hipposideros caffer* in Rwanda (Nziza et al. 2020). Our results again provide some new and consistent data points for similar betacoronavirus diversity and prevalence in East Africa.

In addition to detection of sarbecoviruses in *Hipposideros* and *Rhinolophus* spp. bats, we also detected a sarbecovirus in Angolan rousette bats (*Myonycteris angolensis*). A betacoronavirus was reported from *M. angolensis* (listed as *Rousettus angolensis* in the paper) in Rwanda (Nziza et al. 2020). Betacoronaviruses have also been detected in *Myonycteris torquata* in the Democratic Republic of the Congo (Meta Djomsi et al. 2023) and Cameroon (Ntumvi et al. 2022). Djmosi et al. (Meta Djomsi et al. 2023) found a nobecovirus in *Myonycteris* from west/central Africa; nobecoviruses represented most of the betacoronavirus detections from *Myonycteris* bats in this study (Figures 3 and 4). The strain of sarbecovirus from *Myonycteris* clusters phylogenetically with the strain detected in *Rhinolophus* (Figure 3). These bats have been observed co-roosting in the past (unpublished data) but during this study these bats were inhabiting different caves. Similarly, one individual *Myonycteris* bat was positive for hibecovirus RNA, which we predominantly detected in *Hipposideros* bats (Figures 3, 4). It is unclear how or where cross-species viral sharing occurred, although these two species of bats could have been co-roosting in a cave not sampled during this study. Nziza et al., 2020 detected a new sarbecovirus strain in both *Hipposideros* and *Rhinolophus* co-roosting in a cave. Chidoti et. al., 2022 also reported the presence of 8 groups (strains) of alphacoronaviruses and 8 strains of betacoronaviruses circulating amongst multiple bat species in three study locations. Sequences from each virus group were detected in multiple bat species co-roosting at these locations during a similar time frame, with prevalences ranging from 16 – 39%. The authors postulated that this high diversity of coronaviruses and synchronous periods of viral shedding creates a vulnerability for cross-species transmission and emergence of new strains. Our data provide additional support for there being a high diversity of coronaviruses in these taxa of bats, but that cross-species transmission among co-roosting bats does occur.

Homologous recombination events were detected in some of the sarbecoviruses. While there were clear parental signals consistent with natural recombination events, a known phenomenon in coronaviruses (Simon-Loriere and Holmes 2011), we still cannot rule out the possibility of PCR-mediated recombination during amplification (Yu et al. 2006). Nonetheless, if these homologous recombination events are naturally occurring, they would be consistent with the well-documented evolutionary dynamics of coronaviruses where recombination acts as a major driver for and genetic diversity being facilitated by the tendency of template switching by the viral RNA-dependent RNA polymerase during replication (Lai 1992). Recombination events are particularly common in SARSr-CoVs (sarbecoviruses) and can lead to the emergence of novel viral variants (Boni et al. 2020). For instance, recombination seems to have played a role in the origin of SARS-CoV (Hon et al. 2008) and SARS-CoV-2 (Boni et al. 2020, Lytras et al. 2022).

Our observation of potential recombination events in sarbecoviruses found in Ugandan bats is consistent with findings from other regions in Africa (Woo et al. 2012) and Asia (Lau et al. 2015, Wong et al. 2019). Furthermore, the ecological context in which our study was conducted, that is, the co-roosting of multiple bat genera in the same caves and their harboring of diverse coronaviruses, provides a plausible environment for co-circulation of genetically distinct viral strains, further facilitating recombination. Furthermore, recapture data from PIT tags provided tangible evidence for movement of *Rhinolophus* spp. bats between caves. These findings could be evidence of ongoing evolution of coronaviruses with known zoonotic potential in this region, further reinforcing the significance of continued surveillance in such high human-bat interface regions since recombination can facilitate viral spillovers.

### Paramyxoviruses

Six unique paramyxoviruses were detected over the course of the study; three were from *H.caffer,* one from *N. thebaica,* one from *Rhinolophus* spp., and one from *Myonycteris angolensis*. This viral family is likely underrepresented in our dataset, since we did not collect urine samples where they are most commonly detected (Jolma et al. 2021). The virus detected from *M. angolensis* clusters with the henipaviruses. While some henipaviruses have significant human and veterinary health consequences, such as Nipah and Hendra viruses (Field et al. 2001), the health significance of these strains from African bats is unknown. Henipaviruses have previously been detected from bats in Africa, particularly *Eidolon helvum*; there has also been evidence of spillover to humans and livestock although in the absence of reported outbreaks (Mbu’u et al. 2019, Madera et al. 2022).

Three of the paramyxoviruses detected in *Hipposideros* were similar to previously reported sequences from the PREDICT project (Fig 5). Similarly, the virus detected in this study from *Nycteris thebaica* (bat 279) is also similar to a virus reported from PREDICT (Figure 5). These paramyxovirus strains cluster with the genus *Jeilongvirus*. Jeilongviruses have a global distribution and have been detected primarily in rodents and bats (Zhu et al. 2022, Ch’ng et al. 2023, DeRuyter et al. 2024, Gan et al. 2024). While Jeilongviruses have been shown to replicate in human and non-human primate cells (DeRuyter et al. 2024), there is no evidence that viruses in this genus present a human health threat. Viral RNA from a Jeilong-related virus strain was found in the feces of *Myotis emarginatus* bats, and from *Rhinolophus ferrumequinum* bats co-roosting with *Myotis* in Luxembourg (Pauly et al. 2017). Alpha– and betacoronaviruses were also co-circulating in these bat colonies. Later, a Jeilong-related virus that clusters with the strains from Luxembourg was detected in a *Hipposideros* bat in Zimbabwe (Bourgarel et al. 2018). Jeilongviruses have also been detected in *Hipposideros* bats in China (Zhu et al. 2022) and *Miniopterus schreibersii* in Korea (Noh et al. 2018). Closely related to this strain from Korea are two additional strains isolated from oral swab samples of the Eastern bent-winged bat (*Miniopterus fuliginosus*) and the Far Eastern myotis bat (*Myotis bombinus*) in Japan (Sata et al. 2024). Our data support the widespread distribution of Jeilongviruses in Old World bat taxa.

**Figure 5.**
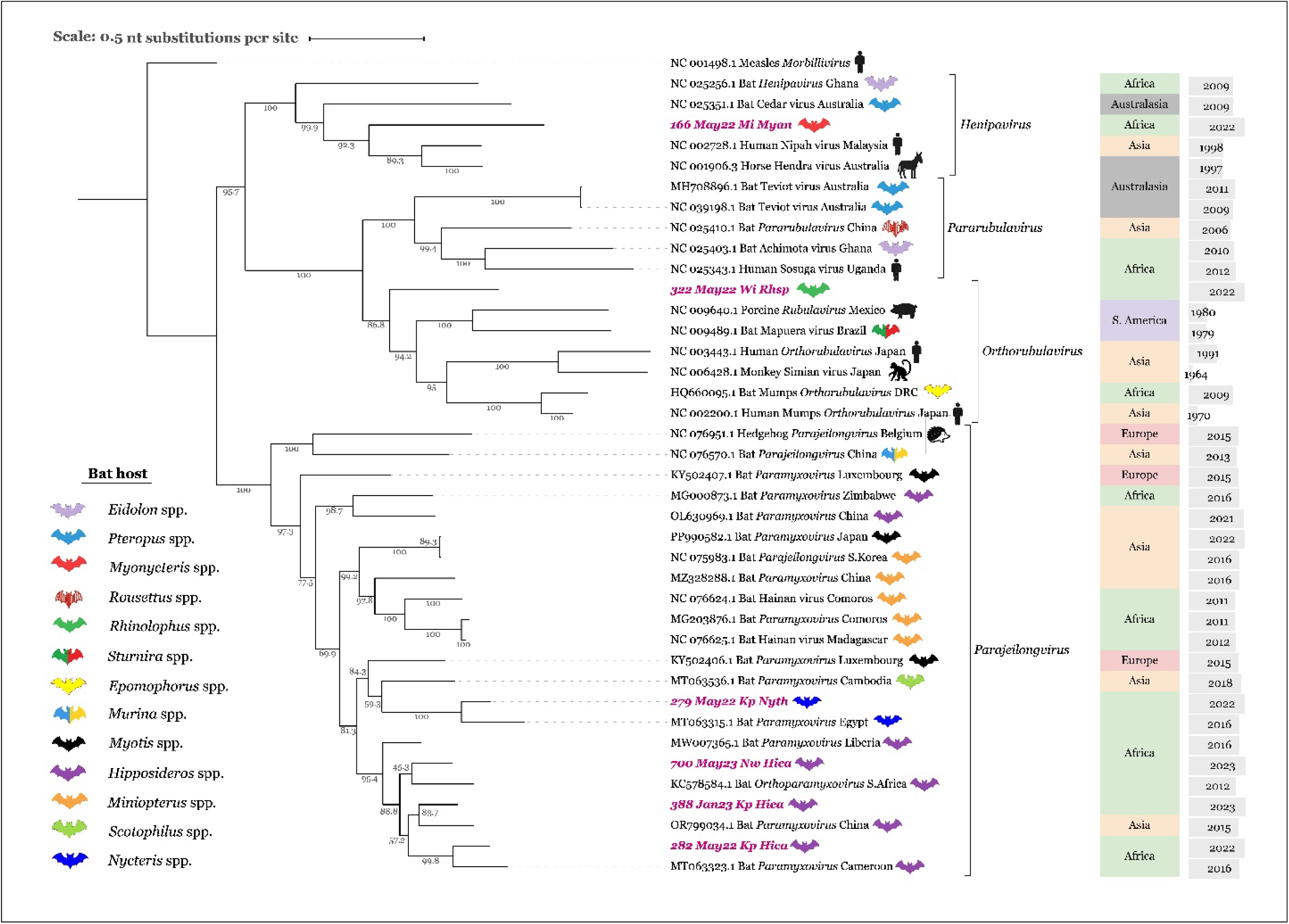
Maximum likelihood tree based on nucleotide alignment of known paramyxoviruses alongside those sequences detected in *Myonycteris, Nycteris*, and *Hipposideros* bats. Aligned sequences represent a conserved fragment of the pol (L) gene (Tong, 2008) Phylogenetic tree rooted by Measles morbillivirus. Sequences detected in current study are highlighted with pink text. Branch support values represent bootstrap percentages based on 1,000 replicates. Icons right of the branch labels represent host species of the virus, with the continent on which the sample was collected and the collection year.

### Rhabdoviruses

In this study we detected three unique rhabdoviruses, all from *Rhinolophus* spp. Of potential importance to animal health was a vesiculovirus in a *Rhinolophus* spp. (Figure 6). Vesiculoviruses are yet another genus of rhabdovirus with a growing body of evidence for association with bats. Vesiculoviruses cause vesiculating (blister-like) lesions in mammals (Sata et al. 2024); many are veterinary pathogens that cause disease in livestock, such as vesicular stomatitis virus which is a widespread disease threat in the Americas (Pelzel-McCluskey 2024). Transmission is recognized to be vector-borne but can also occur by direct contact (Pelzel-McCluskey 2024). Some vesiculoviruses are known to cause febrile disease and outbreaks (Chandipura virus) in people, including veterinarians and animal handlers who have contact with sick animals (Tesh et al. 1977, Basak et al. 2007, Sata et al. 2024). Many vesiculoviruses have also been described from bats (Sata et al. 2024). It is unclear whether these viruses are of concern to human or livestock health, but vesiculoviruses appear to be globally distributed in various insectivorous bat species. Mediterranean bat virus has been described from bats in the genera *Rhinolophus* and *Miniopterus* (Luo et al. 2025). Multiple vesiculoviruses have been reported from bats in China, including Yinshui bat virus from *Rhinolophus* bats (Luo et al. 2021) and Jihong bat virus and Benxi bat virus (Xu et al. 2018). Mejal virus is a related strain sequenced from streblid bat flies collected on Parnell’s mustached bats (*Pteronotus parnellii*) in Mexico (Ramirez-Martinez et al. 2021). Also from the Americas is American bat vesiculovirus in big brown bats (*Eptesicus fuscus*) (Ng et al. 2013).

**Figure 6.**
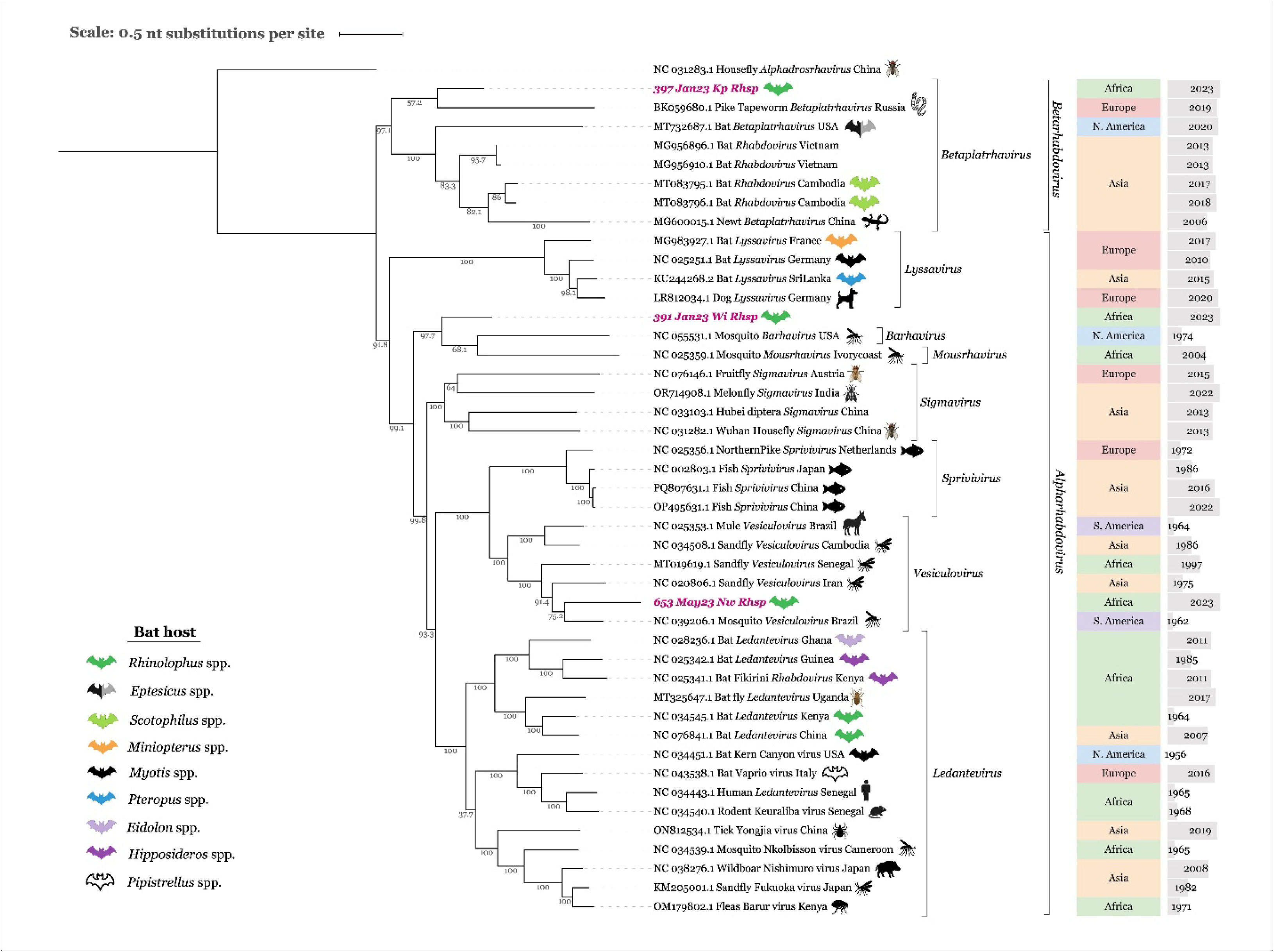
Maximum likelihood tree based on nucleotide sequences of rhabdoviruses. Aligned sequences represent a conserved 260 nt fragment of the rhabdovirus L gene (Balinandi, 2022) The tree is rooted by Wuhan house fly virus 2. Branch support values represent bootstrap percentages based on 1,000 replicates. Icons right of the branch labels represent host species of the virus, with the continent on which the sample was collected and the collection year.

In 1969, Mt. Elgon bat virus (MEBV; *Rhabdoviridae, Ledantevirus*) was first isolated and characterized from a *Rhinolophus* spp. bat captured in the same geographic area in which this study was conducted (Metselaar et al. 1969). A very similar virus was isolated from a roundleaf bat (*Hipposideros vittatus*) in Kenya (Binger et al. 2015) and from straw-colored flying foxes (*Eidolon helvum*) in Ghana (Binger et al. 2015). A number of additional bat-associated viruses clustering phylogenetically close to MEBV led to the formal characterization of the genus *Ledantevirus* (Blasdell et al. 2015). Intriguingly, a multitude of virus strains in the genus *Ledantevirus* were detected in ectoparasitic bat flies (family *Nycteribiidae*) parasitizing Angolan soft-furred bats (*Myonycteris angolensis ruwenzorii*) in western Uganda; viral RNA was detected in the saliva of one bat (Bennett et al. 2020). While we did not detect MEBV or any other ledanteviruses in this study, the presence of these viruses in Uganda but lack of detection in this study is noteworthy.

One of the more unique findings from this study is the detection of a platrhavirus in a *Rhinolophus* spp. Platrhaviruses parasitize vertebrate and invertebrate hosts, and have been sequenced from flatworms (Dheilly et al. 2022). Detection of these viruses in vertebrate hosts, such as in wildlife fecal samples (Bodewes et al. 2014), may indicate a platyhelminth infection (Dheilly et al. 2022). Numerous reports of rhabdoviruses associated with ectoparasites of bats have arisen and been discussed in the context of “viral hyperparasitism” to describe the diversity of microbes that infect parasites of the vertebrate host (Tendu et al. 2022). This hypothesis was also raised by Hause et al., (2020) who detected Sodak rhabdovirus 1 and 2 in the gut viscera of big brown bats in the United States (Hause et al. 2020). Since Sodak rhabdovirus 1 is most likely a virus of nematodes, and Sodak rhabdovirus 2 of insects, this result was interpreted as being from a parasitic infection or dietary content (Hause et al. 2020). The virus detected from Bat 397 clusters with a virus from a tapeworm, Sodak rhabdovirus 2, and similar unclassified viruses that have been previously detected in the fecal samples or oral swabs of bats captured in Cambodia and Vietnam (Figure 6). Our results are consistent with rhabdovirus 397 representing a hyperparasitic virus associated with either a parasite in the bat, or an insect prey species. Parasite infections have been found to be very common in African bat species, and include hemosporidian parasites in the genera *Plasmodium*, *Polychromophilus*, *Nycteria*, and *Hepatocystis* (Schaer et al. 2013), trypanosomes (Thiombiano et al. 2023), and filarial nemotodes in the genus *Litmosa* (Pikula et al. 2023). In Cameroon, parasites in the genera *Nycteria* ranged in prevalence from around 20% in *Rhinolophus landeri* bats to 60-65% in other bat genera (Martin et al. 2006). Approximately 36% of *R. alcyone* sampled in Burkina Faso were infected with trypanosomes (Thiombiano et al. 2023). One *Litmosa* nematode species was first described from *R. ferrumequinum* from Algeria (Martin et al. 2006). Therefore, platrhavirus infection of a parasite in the bat is plausible, and could explain other published, uncharacterized rhabdovirus sequences from bat fecal samples (i.e. GenBank accessions QID58057, QID58043, URC21140, URC21139). Bat endoparasites and the viruses they may be infected with are not well characterized and represent an area of future investigation with lots of discovery potential.

## Conclusions

Over the course of conducting viral surveillance of bats in Eastern Uganda, we detected diverse coronaviruses, paramyxoviruses, and rhabdoviruses. The strains detected in this study from each viral family represent strains related to those reported previously in other parts of Africa, but provide additional geographic context for virus-bat associations. Evidence for viral sharing was noted among the bats and coronaviruses, with spillover of a minunacovirus from *Miniopterus* to *Hipposideros*, a nobecovirus from *Myonycteris* to *Rhinolophus*, a hibecovirus from *Hipposideros* to *Myonycteris*, and a sarbecovirus from *Rhinolophus* to *Myonycteris* and *Hipposideros*. These results are consistent with other reports of coronavirus circulation in African bats and have implications for virus evolution and the recognized diversity of viruses in this family. The health significance of any of the virus strains detected in this study to people or animals is unknown. Some viral strains were related to those known to cause disease in humans (coronaviruses, henipaviruses) or animals (vesiculoviruses) while others seem to be strains that naturally circulate in wildlife (jeilongviruses, platraviruses) with no known public health significance. In general, rhabdoviruses and paramyxoviruses are commonly detected in bats, but many strains are uncharacterized and unclassified. A deeper understanding of the viral ecology for strains with potential health consequences, and assessment of human exposure risk in the cave environment is warranted.

## Acknowledgements

We sincerely thank the people of the Kapchorwa and Sipi towns who granted permission for this sampling to occur. In particular, we thank Dennis Ssemwogere and Godfrey Kyazze for transportation, Winnex Cherotwo for facilitating community engagements, as well as Enock Chelangat, Millat Tongo, Prisca Chekwemboi; Junia Chebet, and Phalti Chebet.

## Ethical Approvals

All applicable institutional and/or national guidelines for the care and use of animals were followed. Field sampling of bats was approved by the Uganda Wildlife Authority (COD/96/05), the Uganda National Council for Science and Technology (NS663), Institutional Animal Care and Use Committee protocol 1314, the US Department of Defense Animal Care and Use Committee (ACURO)(CT-2019-21.e001). All applicable institutional and/or national guidelines for the care and use of animals were followed.

## Sample importation

Importation of bat-derived samples was conducted under CDC PHS permits 20200922-3028A (2021-2022), 20220613-2125A (2022-2023), and 20230610-2285A (2023-2024) and USFWS Dec Control Num: 2022-109695; 2023-285124.

## Biosafety

Sample processing at Colorado State University was approved by the CSU Institutional Biosafety Committee (18-083B).

**Supplementary Table 1:**
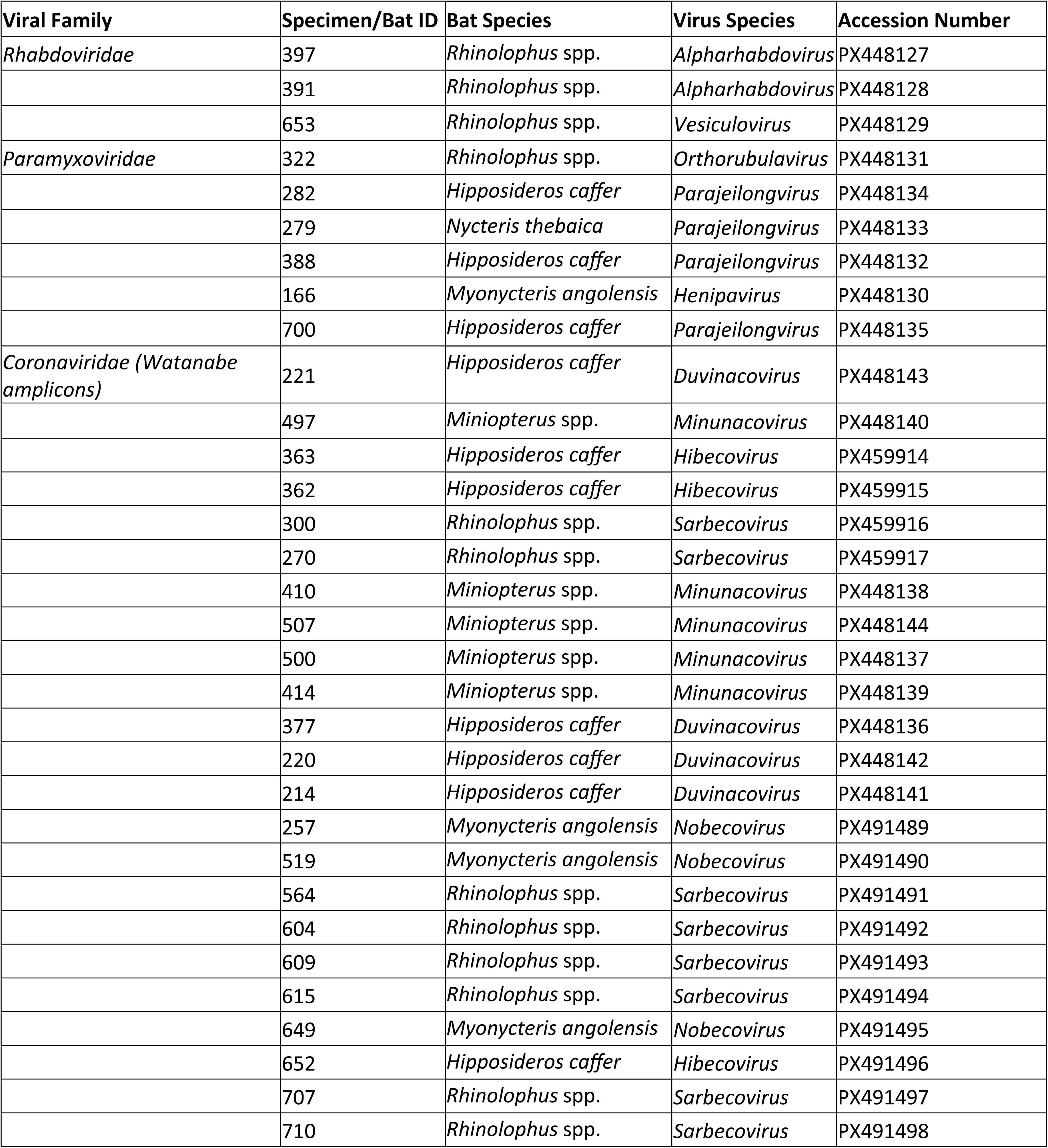
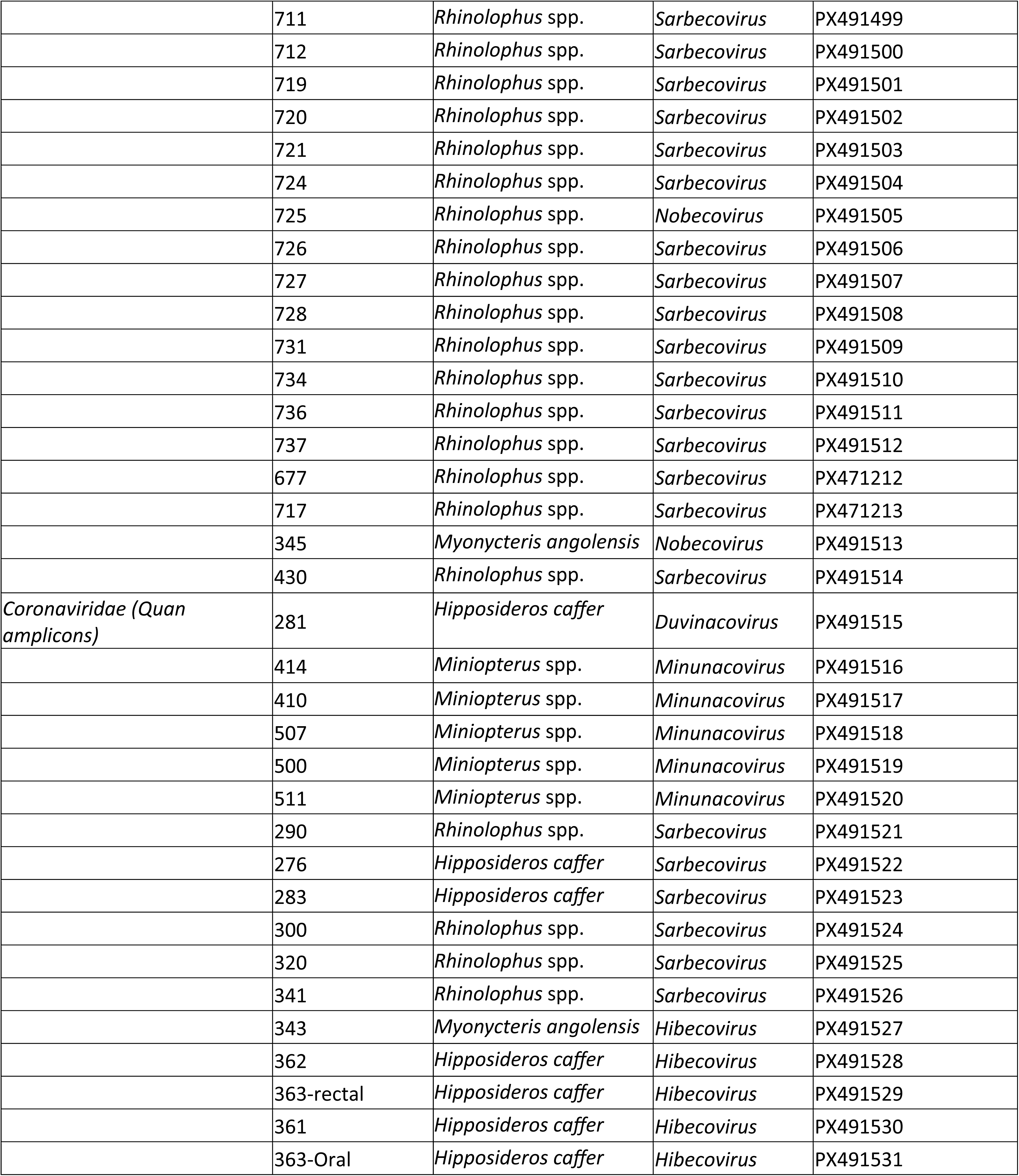
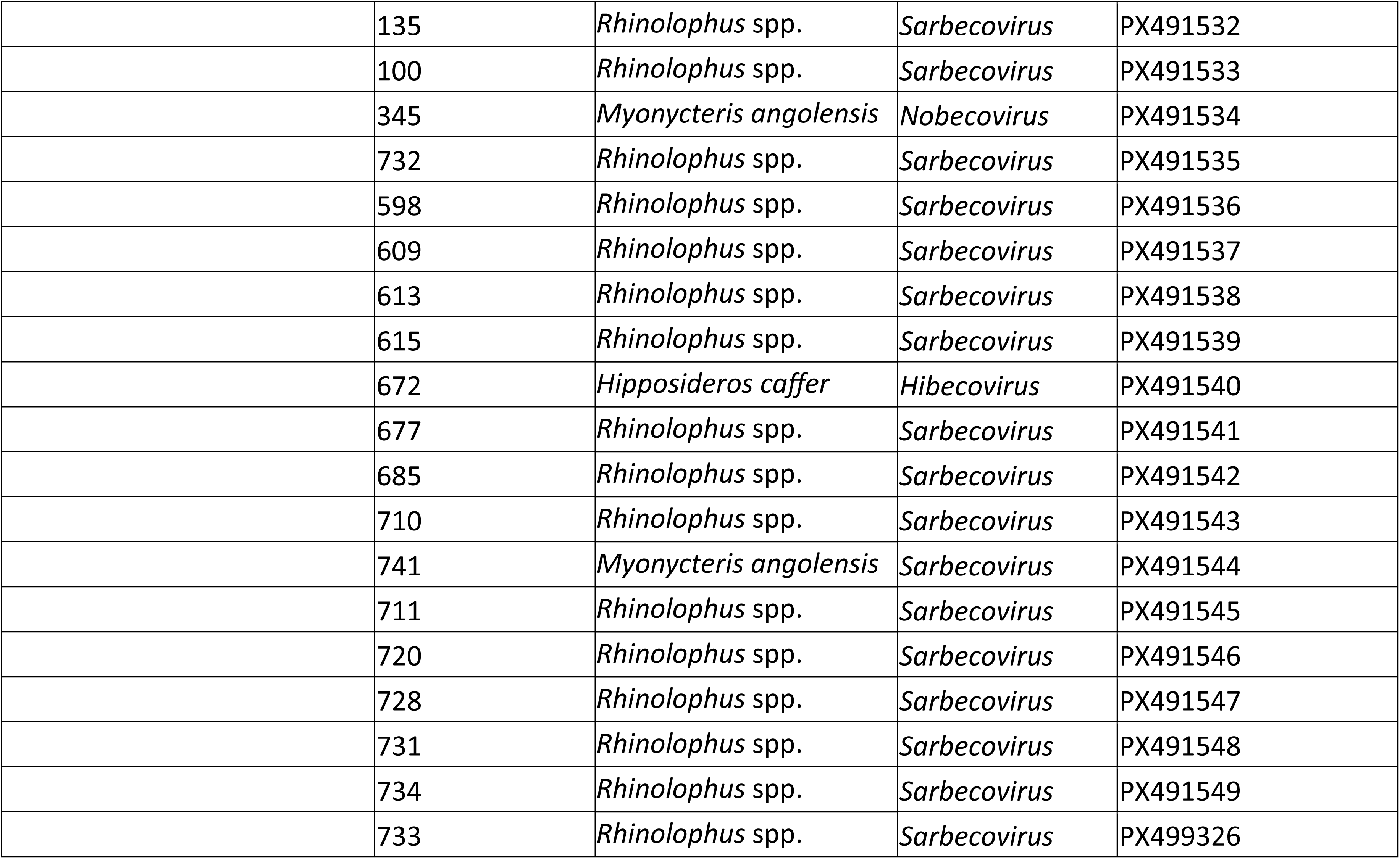
GenBank Accession Numbers.

**Supplementary Table 2:**
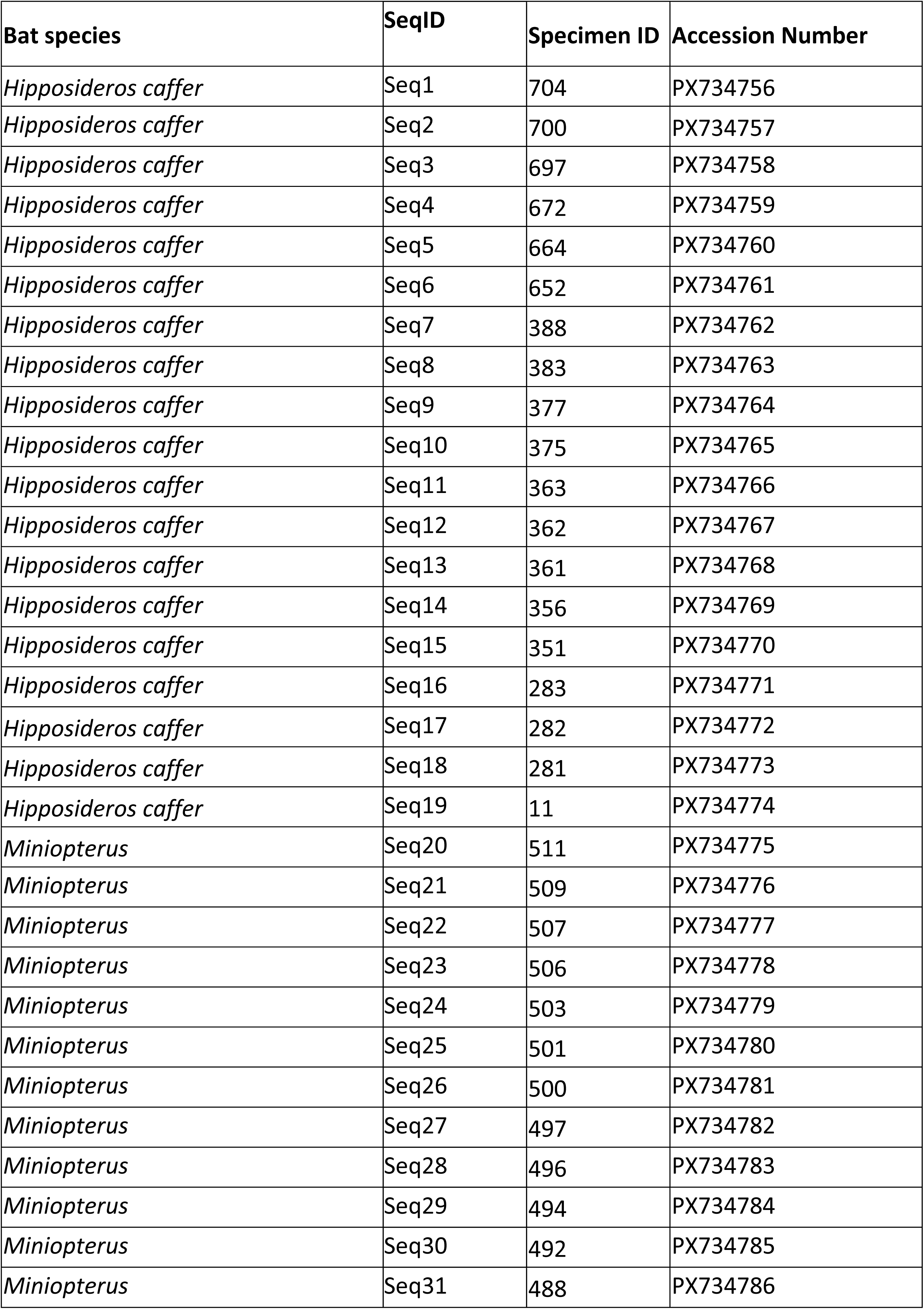
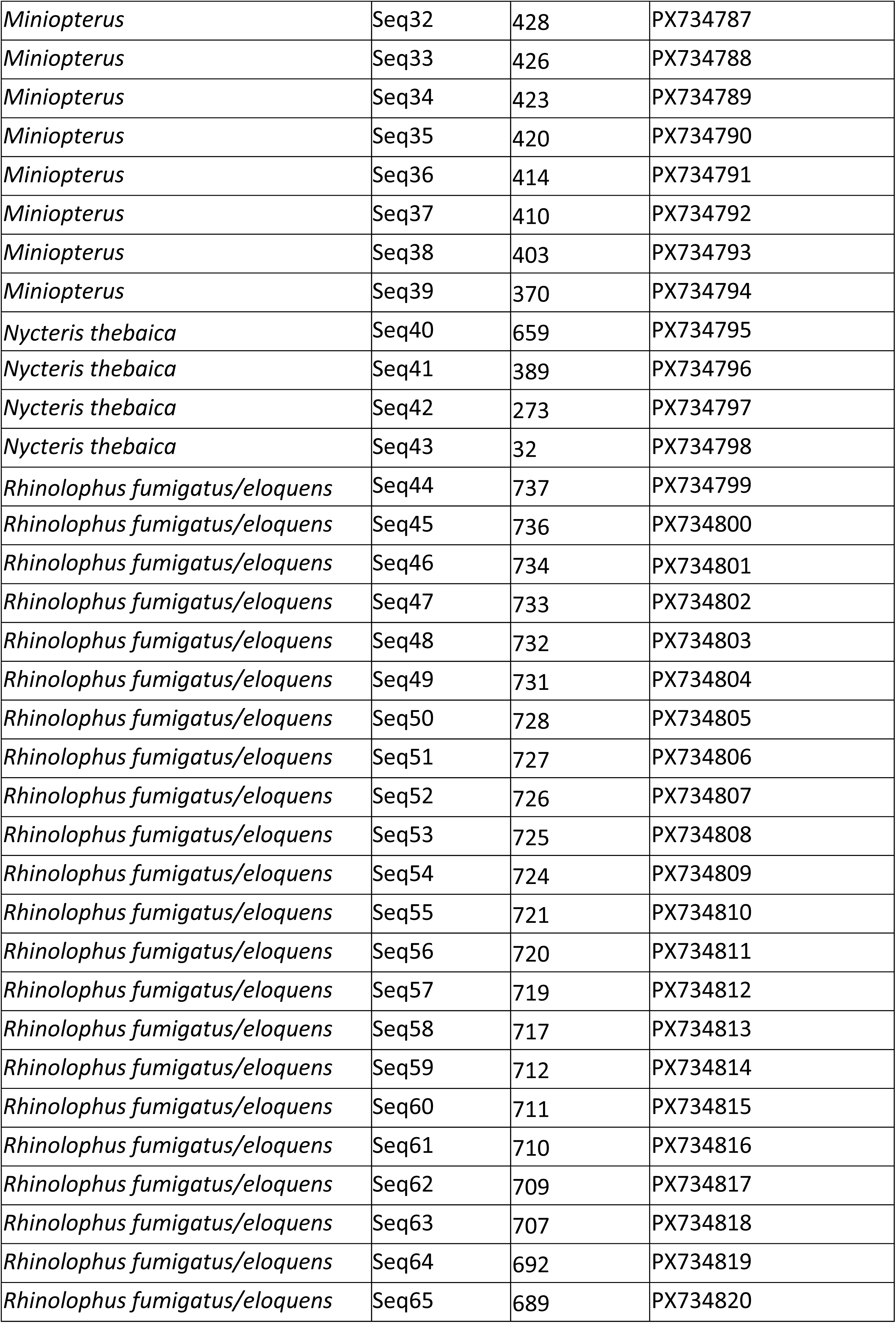
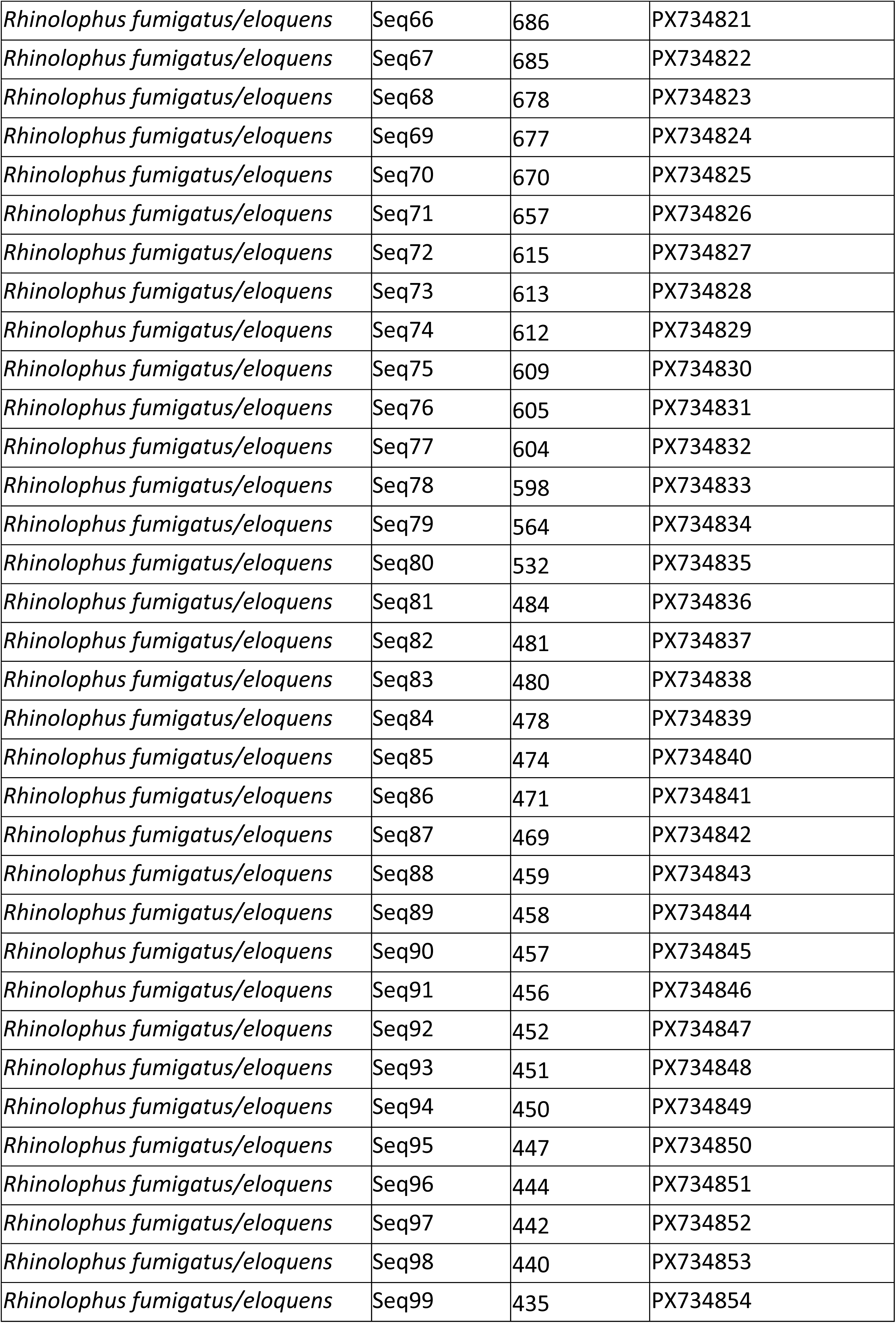
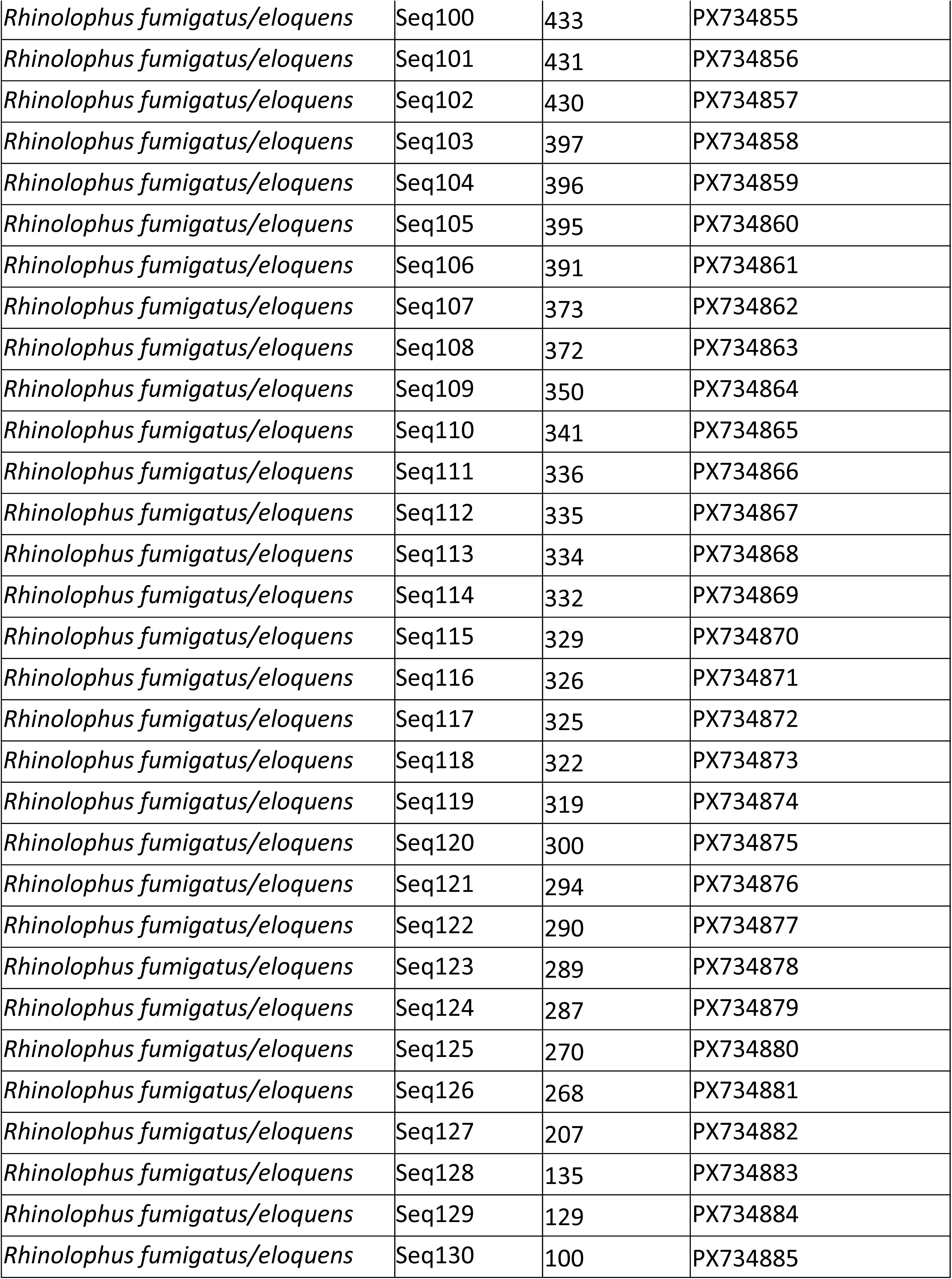
GenBank Accession Numbers for bat cytochrome b sequences.

| Bat species | SeqID | Specimen ID | Accession Number |
| --- | --- | --- | --- |
| *Hipposideros caffer* | Seq1 | 704 | PX734756 |
| *Hipposideros caffer* | Seq2 | 700 | PX734757 |
| *Hipposideros caffer* | Seq3 | 697 | PX734758 |
| *Hipposideros caffer* | Seq4 | 672 | PX734759 |
| *Hipposideros caffer* | Seq5 | 664 | PX734760 |
| *Hipposideros caffer* | Seq6 | 652 | PX734761 |
| *Hipposideros caffer* | Seq7 | 388 | PX734762 |
| *Hipposideros caffer* | Seq8 | 383 | PX734763 |
| *Hipposideros caffer* | Seq9 | 377 | PX734764 |
| *Hipposideros caffer* | Seq10 | 375 | PX734765 |
| *Hipposideros caffer* | Seq11 | 363 | PX734766 |
| *Hipposideros caffer* | Seq12 | 362 | PX734767 |
| *Hipposideros caffer* | Seq13 | 361 | PX734768 |
| *Hipposideros caffer* | Seq14 | 356 | PX734769 |
| *Hipposideros caffer* | Seq15 | 351 | PX734770 |
| *Hipposideros caffer* | Seq16 | 283 | PX734771 |
| *Hipposideros caffer* | Seq17 | 282 | PX734772 |
| *Hipposideros caffer* | Seq18 | 281 | PX734773 |
| *Hipposideros caffer* | Seq19 | 11 | PX734774 |
| *Miniopterus* | Seq20 | 511 | PX734775 |
| *Miniopterus* | Seq21 | 509 | PX734776 |
| *Miniopterus* | Seq22 | 507 | PX734777 |
| *Miniopterus* | Seq23 | 506 | PX734778 |
| *Miniopterus* | Seq24 | 503 | PX734779 |
| *Miniopterus* | Seq25 | 501 | PX734780 |
| *Miniopterus* | Seq26 | 500 | PX734781 |
| *Miniopterus* | Seq27 | 497 | PX734782 |
| *Miniopterus* | Seq28 | 496 | PX734783 |
| *Miniopterus* | Seq29 | 494 | PX734784 |
| *Miniopterus* | Seq30 | 492 | PX734785 |
| *Miniopterus* | Seq31 | 488 | PX734786 |
| *Miniopterus* | Seq32 | 428 | PX734787 |
| *Miniopterus* | Seq33 | 426 | PX734788 |
| *Miniopterus* | Seq34 | 423 | PX734789 |
| *Miniopterus* | Seq35 | 420 | PX734790 |
| *Miniopterus* | Seq36 | 414 | PX734791 |
| *Miniopterus* | Seq37 | 410 | PX734792 |
| *Miniopterus* | Seq38 | 403 | PX734793 |
| *Miniopterus* | Seq39 | 370 | PX734794 |
| *Nycteris thebaica* | Seq40 | 659 | PX734795 |
| *Nycteris thebaica* | Seq41 | 389 | PX734796 |
| *Nycteris thebaica* | Seq42 | 273 | PX734797 |
| *Nycteris thebaica* | Seq43 | 32 | PX734798 |
| *Rhinolophus fumigatus/eloquens* | Seq44 | 737 | PX734799 |
| *Rhinolophus fumigatus/eloquens* | Seq45 | 736 | PX734800 |
| *Rhinolophus fumigatus/eloquens* | Seq46 | 734 | PX734801 |
| *Rhinolophus fumigatus/eloquens* | Seq47 | 733 | PX734802 |
| *Rhinolophus fumigatus/eloquens* | Seq48 | 732 | PX734803 |
| *Rhinolophus fumigatus/eloquens* | Seq49 | 731 | PX734804 |
| *Rhinolophus fumigatus/eloquens* | Seq50 | 728 | PX734805 |
| *Rhinolophus fumigatus/eloquens* | Seq51 | 727 | PX734806 |
| *Rhinolophus fumigatus/eloquens* | Seq52 | 726 | PX734807 |
| *Rhinolophus fumigatus/eloquens* | Seq53 | 725 | PX734808 |
| *Rhinolophus fumigatus/eloquens* | Seq54 | 724 | PX734809 |
| *Rhinolophus fumigatus/eloquens* | Seq55 | 721 | PX734810 |
| *Rhinolophus fumigatus/eloquens* | Seq56 | 720 | PX734811 |
| *Rhinolophus fumigatus/eloquens* | Seq57 | 719 | PX734812 |
| *Rhinolophus fumigatus/eloquens* | Seq58 | 717 | PX734813 |
| *Rhinolophus fumigatus/eloquens* | Seq59 | 712 | PX734814 |
| *Rhinolophus fumigatus/eloquens* | Seq60 | 711 | PX734815 |
| *Rhinolophus fumigatus/eloquens* | Seq61 | 710 | PX734816 |
| *Rhinolophus fumigatus/eloquens* | Seq62 | 709 | PX734817 |
| *Rhinolophus fumigatus/eloquens* | Seq63 | 707 | PX734818 |
| *Rhinolophus fumigatus/eloquens* | Seq64 | 692 | PX734819 |
| *Rhinolophus fumigatus/eloquens* | Seq65 | 689 | PX734820 |
| *Rhinolophus fumigatus/eloquens* | Seq66 | 686 | PX734821 |
| *Rhinolophus fumigatus/eloquens* | Seq67 | 685 | PX734822 |
| *Rhinolophus fumigatus/eloquens* | Seq68 | 678 | PX734823 |
| *Rhinolophus fumigatus/eloquens* | Seq69 | 677 | PX734824 |
| *Rhinolophus fumigatus/eloquens* | Seq70 | 670 | PX734825 |
| *Rhinolophus fumigatus/eloquens* | Seq71 | 657 | PX734826 |
| *Rhinolophus fumigatus/eloquens* | Seq72 | 615 | PX734827 |
| *Rhinolophus fumigatus/eloquens* | Seq73 | 613 | PX734828 |
| *Rhinolophus fumigatus/eloquens* | Seq74 | 612 | PX734829 |
| *Rhinolophus fumigatus/eloquens* | Seq75 | 609 | PX734830 |
| *Rhinolophus fumigatus/eloquens* | Seq76 | 605 | PX734831 |
| *Rhinolophus fumigatus/eloquens* | Seq77 | 604 | PX734832 |
| *Rhinolophus fumigatus/eloquens* | Seq78 | 598 | PX734833 |
| *Rhinolophus fumigatus/eloquens* | Seq79 | 564 | PX734834 |
| *Rhinolophus fumigatus/eloquens* | Seq80 | 532 | PX734835 |
| *Rhinolophus fumigatus/eloquens* | Seq81 | 484 | PX734836 |
| *Rhinolophus fumigatus/eloquens* | Seq82 | 481 | PX734837 |
| *Rhinolophus fumigatus/eloquens* | Seq83 | 480 | PX734838 |
| *Rhinolophus fumigatus/eloquens* | Seq84 | 478 | PX734839 |
| *Rhinolophus fumigatus/eloquens* | Seq85 | 474 | PX734840 |
| *Rhinolophus fumigatus/eloquens* | Seq86 | 471 | PX734841 |
| *Rhinolophus fumigatus/eloquens* | Seq87 | 469 | PX734842 |
| *Rhinolophus fumigatus/eloquens* | Seq88 | 459 | PX734843 |
| *Rhinolophus fumigatus/eloquens* | Seq89 | 458 | PX734844 |
| *Rhinolophus fumigatus/eloquens* | Seq90 | 457 | PX734845 |
| *Rhinolophus fumigatus/eloquens* | Seq91 | 456 | PX734846 |
| *Rhinolophus fumigatus/eloquens* | Seq92 | 452 | PX734847 |
| *Rhinolophus fumigatus/eloquens* | Seq93 | 451 | PX734848 |
| *Rhinolophus fumigatus/eloquens* | Seq94 | 450 | PX734849 |
| *Rhinolophus fumigatus/eloquens* | Seq95 | 447 | PX734850 |
| *Rhinolophus fumigatus/eloquens* | Seq96 | 444 | PX734851 |
| *Rhinolophus fumigatus/eloquens* | Seq97 | 442 | PX734852 |
| *Rhinolophus fumigatus/eloquens* | Seq98 | 440 | PX734853 |
| *Rhinolophus fumigatus/eloquens* | Seq99 | 435 | PX734854 |
| *Rhinolophus fumigatus/eloquens* | Seq100 | 433 | PX734855 |
| *Rhinolophus fumigatus/eloquens* | Seq101 | 431 | PX734856 |
| *Rhinolophus fumigatus/eloquens* | Seq102 | 430 | PX734857 |
| *Rhinolophus fumigatus/eloquens* | Seq103 | 397 | PX734858 |
| *Rhinolophus fumigatus/eloquens* | Seq104 | 396 | PX734859 |
| *Rhinolophus fumigatus/eloquens* | Seq105 | 395 | PX734860 |
| *Rhinolophus fumigatus/eloquens* | Seq106 | 391 | PX734861 |
| *Rhinolophus fumigatus/eloquens* | Seq107 | 373 | PX734862 |
| *Rhinolophus fumigatus/eloquens* | Seq108 | 372 | PX734863 |
| *Rhinolophus fumigatus/eloquens* | Seq109 | 350 | PX734864 |
| *Rhinolophus fumigatus/eloquens* | Seq110 | 341 | PX734865 |
| *Rhinolophus fumigatus/eloquens* | Seq111 | 336 | PX734866 |
| *Rhinolophus fumigatus/eloquens* | Seq112 | 335 | PX734867 |
| *Rhinolophus fumigatus/eloquens* | Seq113 | 334 | PX734868 |
| *Rhinolophus fumigatus/eloquens* | Seq114 | 332 | PX734869 |
| *Rhinolophus fumigatus/eloquens* | Seq115 | 329 | PX734870 |
| *Rhinolophus fumigatus/eloquens* | Seq116 | 326 | PX734871 |
| *Rhinolophus fumigatus/eloquens* | Seq117 | 325 | PX734872 |
| *Rhinolophus fumigatus/eloquens* | Seq118 | 322 | PX734873 |
| *Rhinolophus fumigatus/eloquens* | Seq119 | 319 | PX734874 |
| *Rhinolophus fumigatus/eloquens* | Seq120 | 300 | PX734875 |
| *Rhinolophus fumigatus/eloquens* | Seq121 | 294 | PX734876 |
| *Rhinolophus fumigatus/eloquens* | Seq122 | 290 | PX734877 |
| *Rhinolophus fumigatus/eloquens* | Seq123 | 289 | PX734878 |
| *Rhinolophus fumigatus/eloquens* | Seq124 | 287 | PX734879 |
| *Rhinolophus fumigatus/eloquens* | Seq125 | 270 | PX734880 |
| *Rhinolophus fumigatus/eloquens* | Seq126 | 268 | PX734881 |
| *Rhinolophus fumigatus/eloquens* | Seq127 | 207 | PX734882 |
| *Rhinolophus fumigatus/eloquens* | Seq128 | 135 | PX734883 |
| *Rhinolophus fumigatus/eloquens* | Seq129 | 129 | PX734884 |
| *Rhinolophus fumigatus/eloquens* | Seq130 | 100 | PX734885 |

**Figure S1.**
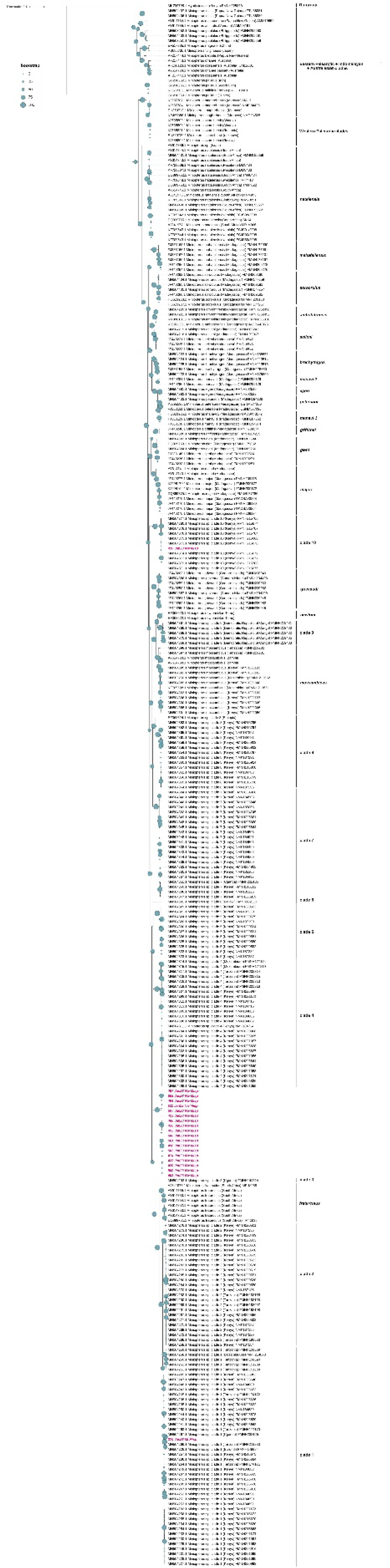
Maximum likelihood tree based on nucleotide alignment of cytochrome b sequences from *Miniopterus* spp. bats, using *Myotis tricolor* as an outgroup. Sequences detected in current study are highlighted with pink text. Branch support values represent bootstrap percentages based on 1,000 replicates; the strength of bootstrap values is indicated by the size of the circle.

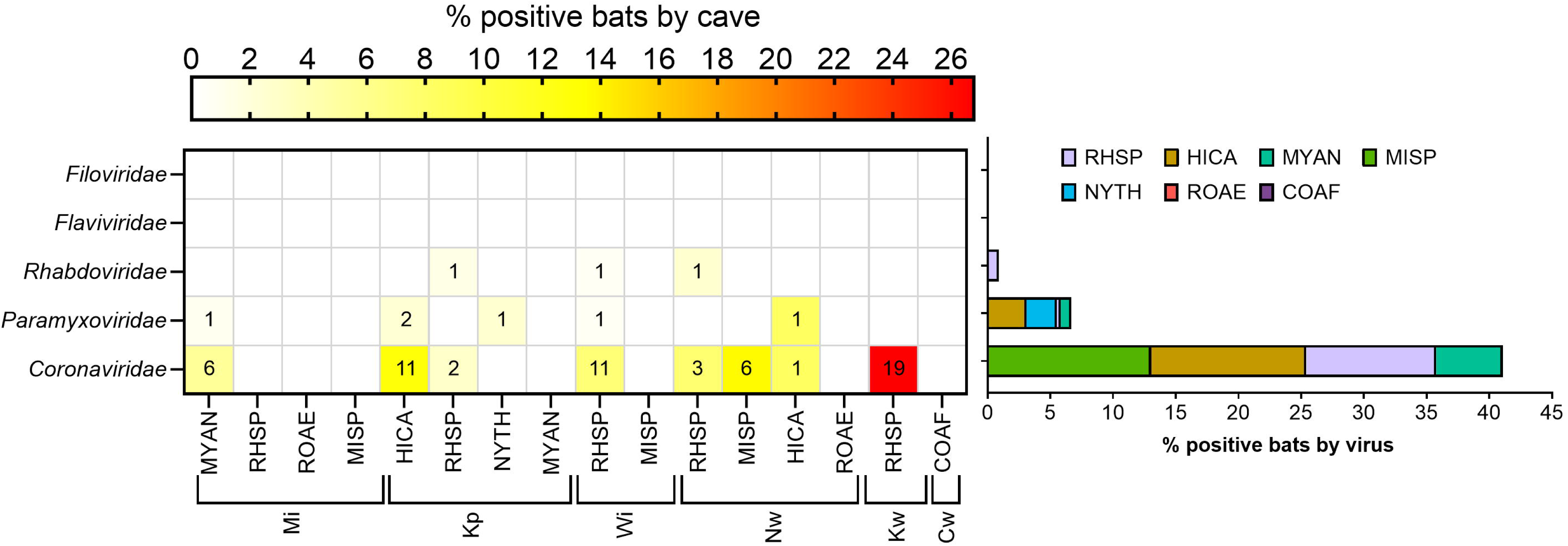

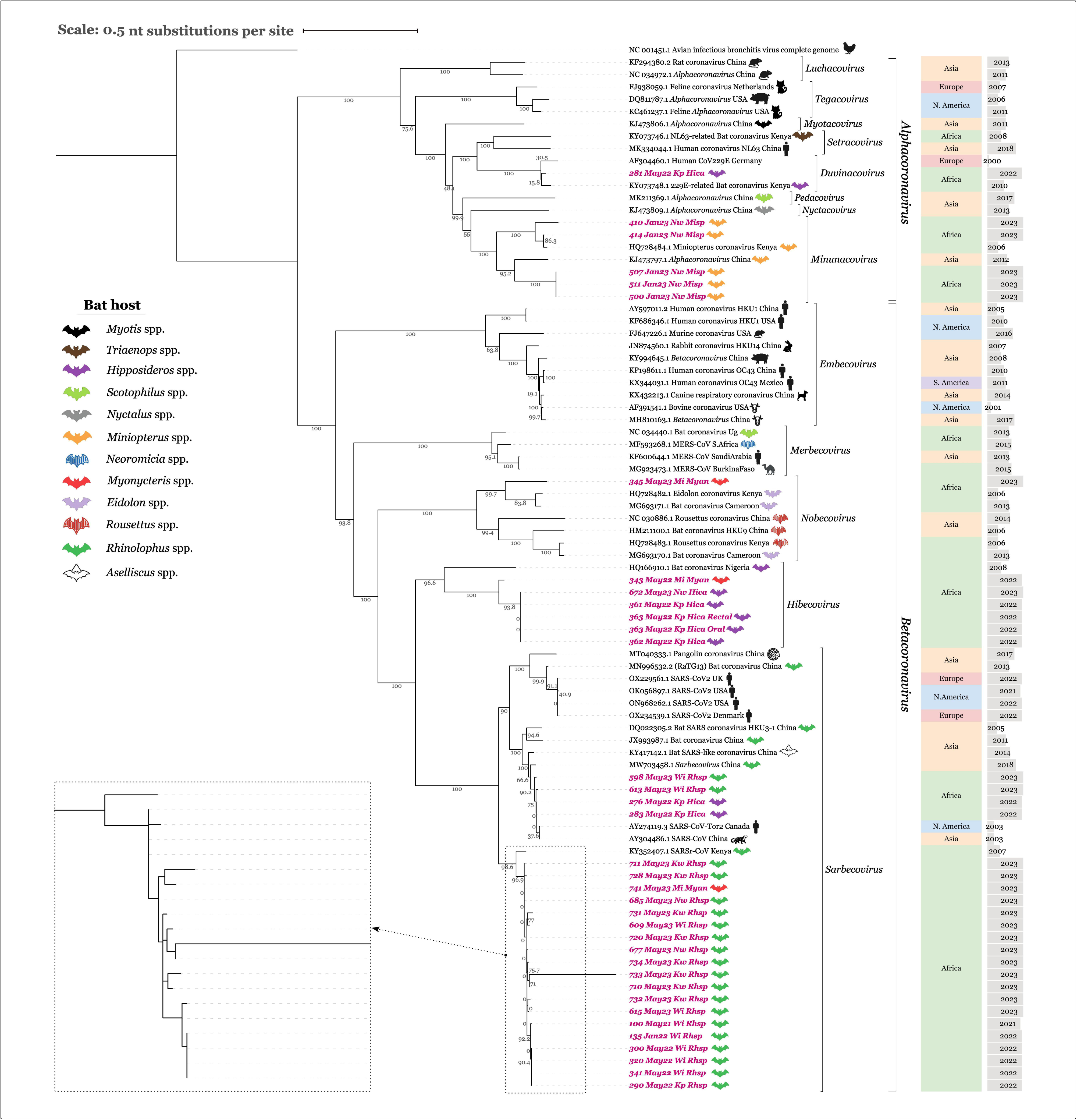

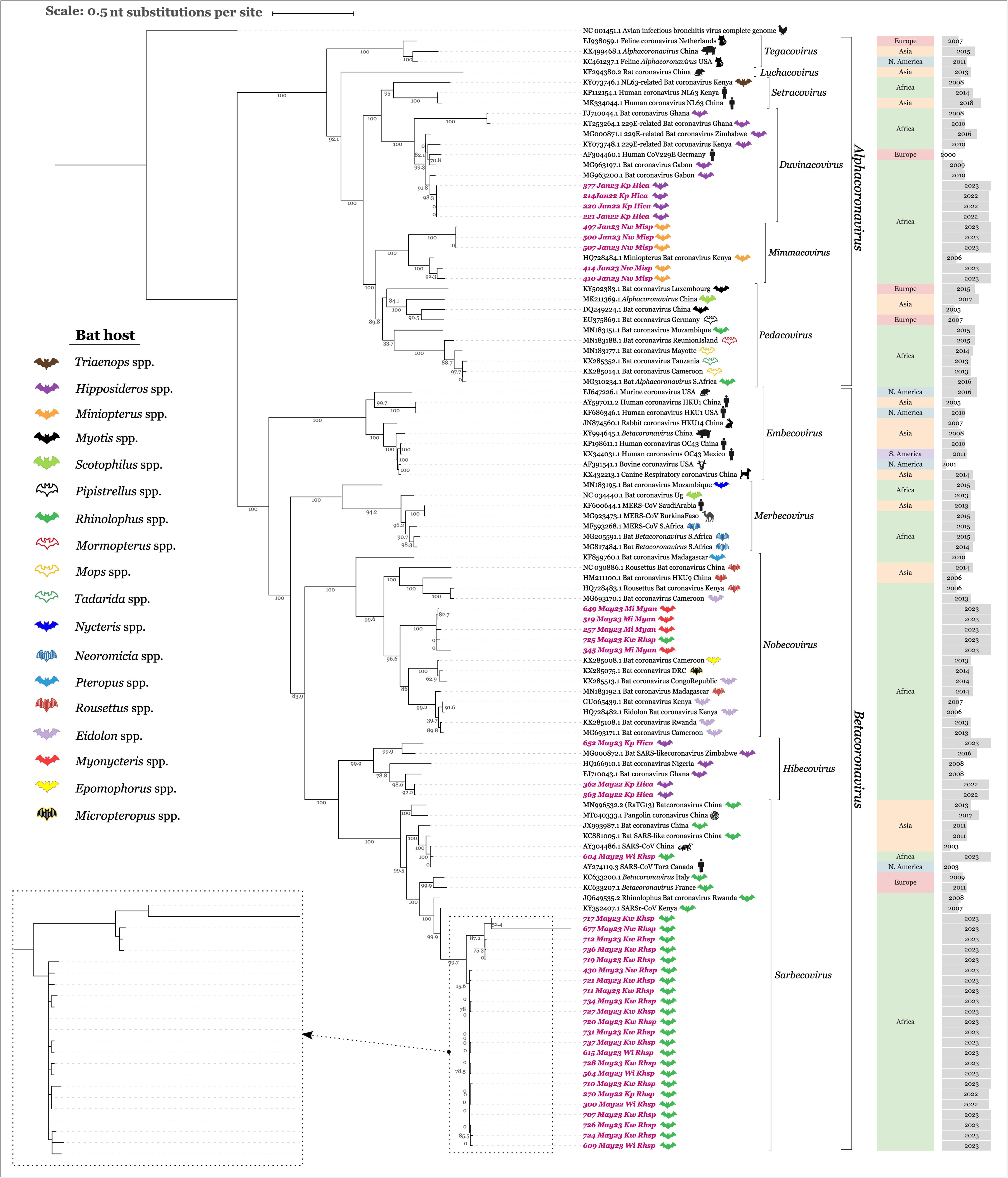

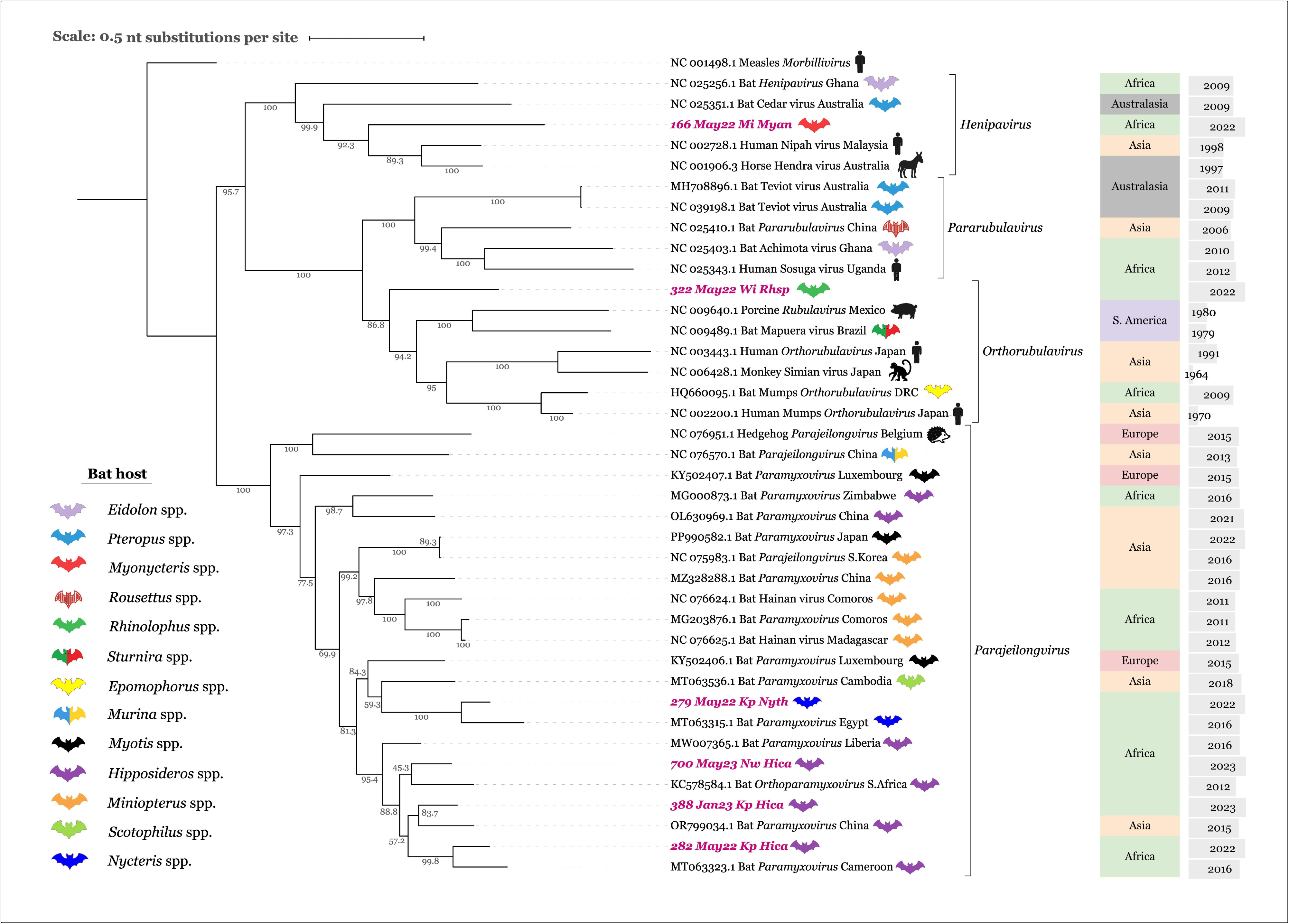

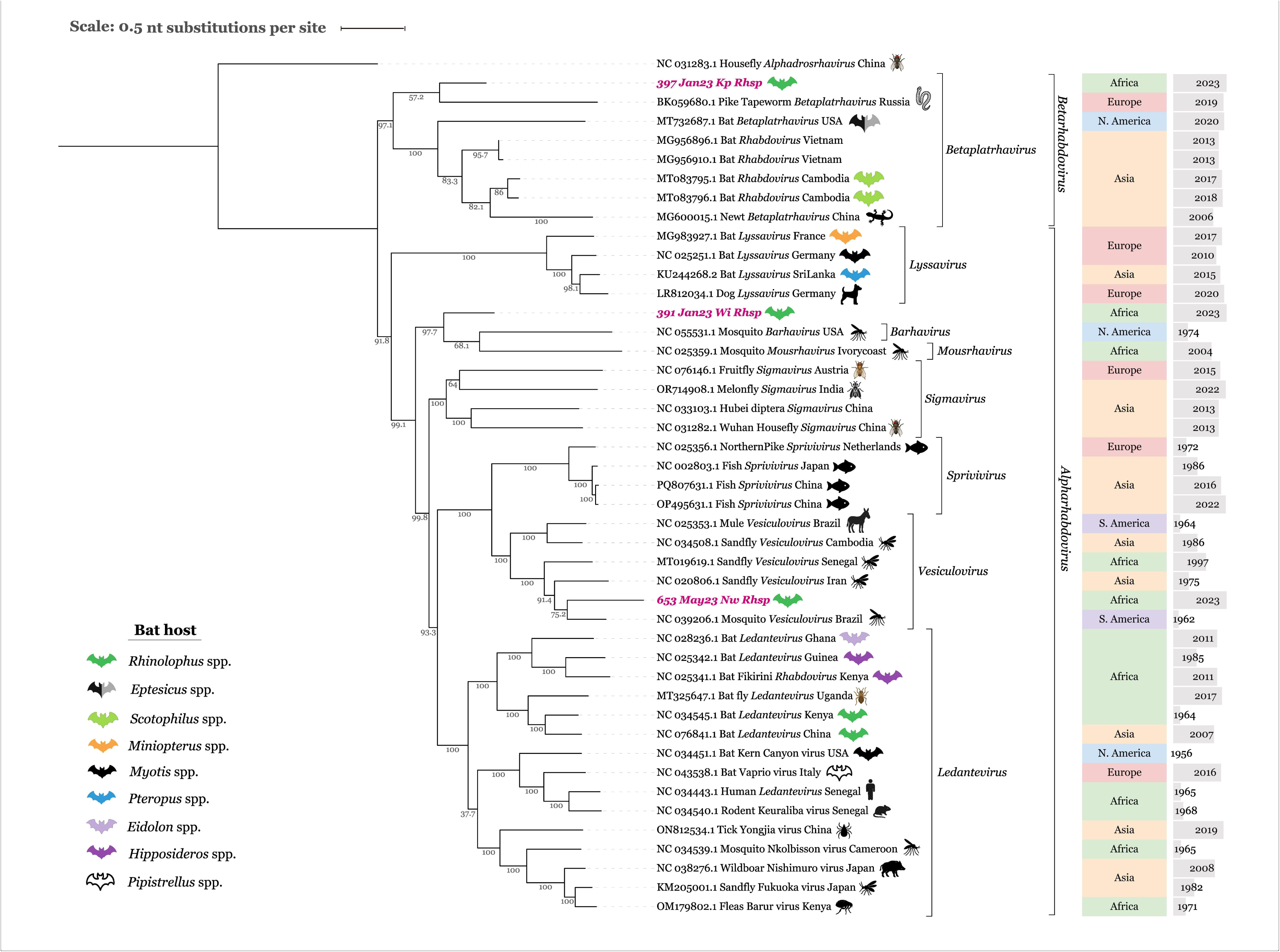

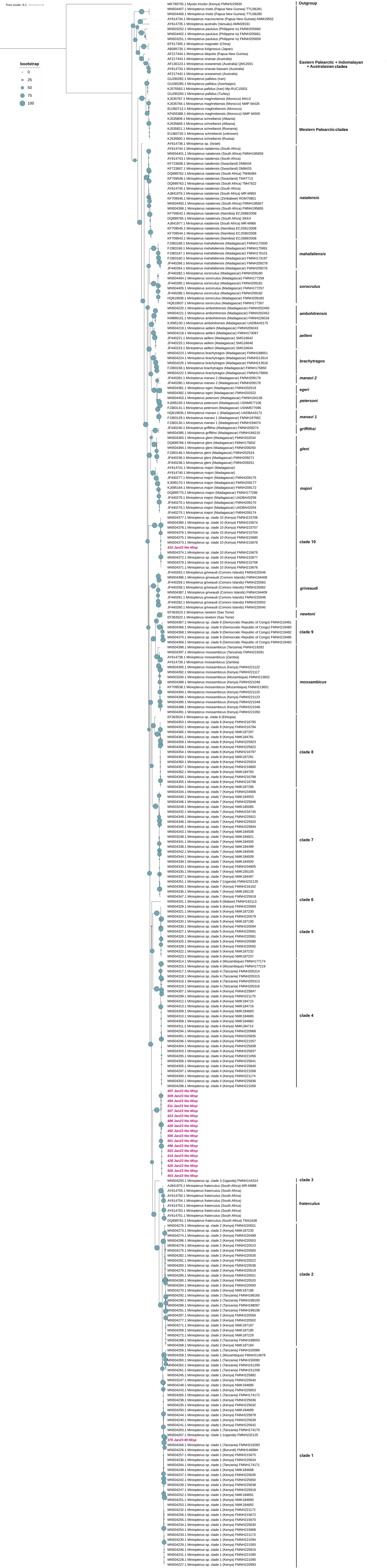

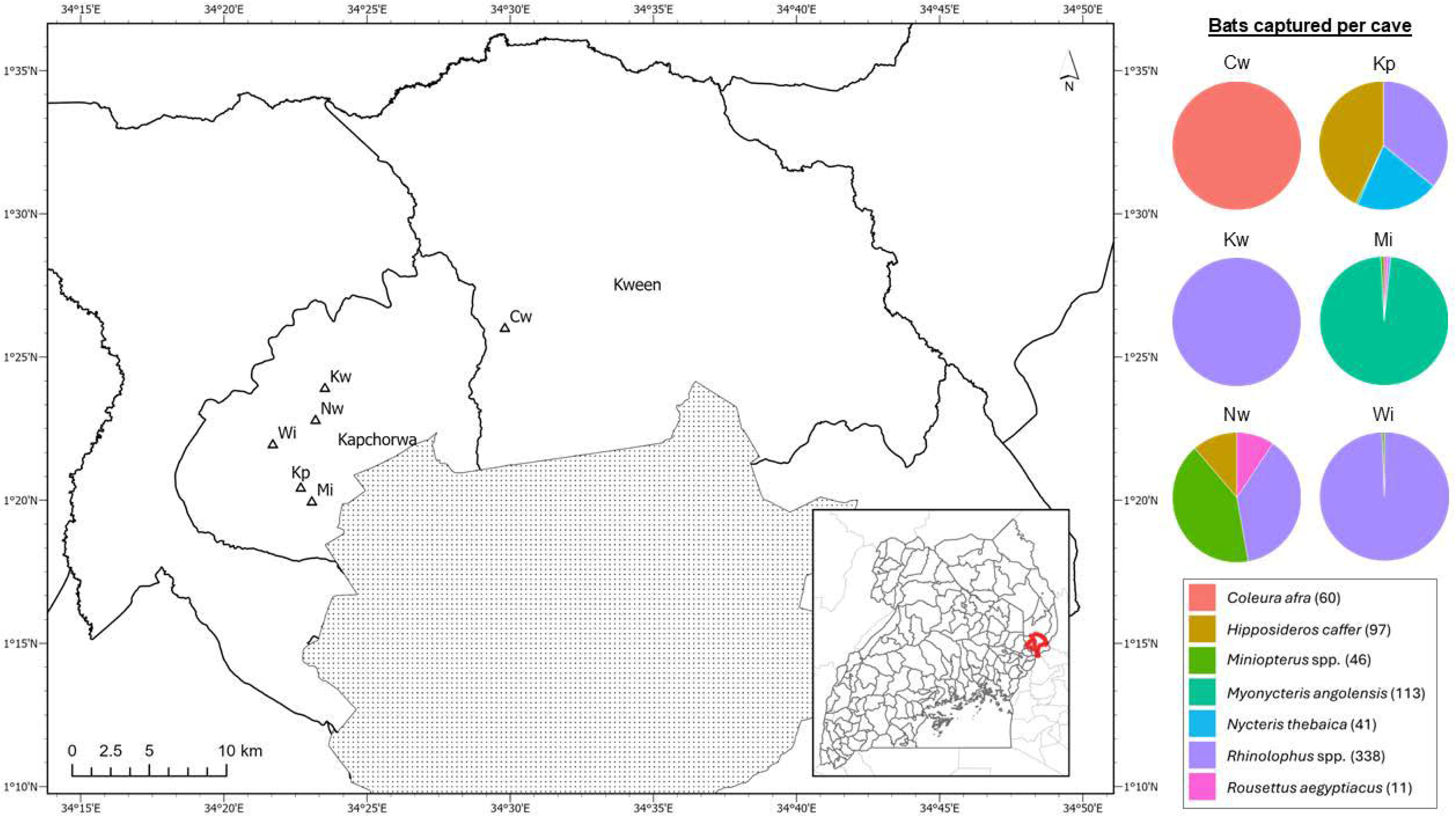

